# High-resolution Linkage Map with Allele Dosage Allows the Identification of an Apomixis Region and Complex Traits in Guinea Grass (*Megathyrsus maximus*)

**DOI:** 10.1101/801399

**Authors:** Thamiris G. Deo, Rebecca C. U. Ferreira, Leticia A. C. Lara, Aline C. L. Moraes, Alessandro Alves-Pereira, Fernanda A. de Oliveira, Antonio A. F. Garcia, Mateus F. Santos, Liana Jank, Anete P. Souza

**Affiliations:** Center for Molecular Biology and Genetic Engineering, University of Campinas, Campinas, Brazil; Genetics Department, Escola Superior de Agricultura “Luiz de Queiroz”, University of Sao Paulo, Piracicaba, Brazil; Embrapa Beef Cattle, Brazilian Agricultural Research Corporation, Campo Grande, Brazil; Plant Biology Department, Biology Institute, University of Campinas, Campinas, Brazil

**Keywords:** apospory, double reduction, forage, polyploidy, QTL, SNP, trait correlations

## Abstract

Forage grasses are mainly used in animal feed to fatten cattle and dairy herds. Among tropical forage crops that reproduce by seeds, guinea grass (*Megathyrsus maximus*) is considered one of the most productive. This species has several genomic complexities, such as autotetraploidy and apomixis, due to the process of domestication. Consequently, approaches that relate phenotypic and genotypic data are incipient. In this context, we built a linkage map with allele dosage and generated novel information about the genetic architecture of traits that are important for the breeding of *M. maximus*. From a full-sib progeny, a linkage map containing 858 single nucleotide polymorphism (SNP) markers with allele dosage information expected for an autotetraploid was obtained. The high genetic variability of the progeny allowed us to map ten quantitative trait loci (QTLs) related to agronomic traits, such as regrowth capacity and total dry matter, and 36 QTLs related to nutritional quality, which were distributed among all homology groups (HGs). Various overlapping regions associated with the quantitative traits suggested QTL hotspots. In addition, we were able to map one locus that controls apospory (apo-locus) in HG II. A total of 55 different gene families involved in cellular metabolism and plant growth were identified from markers adjacent to the QTLs and apomixis locus by using the *Panicum virgatum* genome as a reference in comparisons with the genomes of *Arabidopsis thaliana* and *Oryza sativa*. Our results provide a better understanding of the genetic basis of reproduction by apomixis and traits important for breeding programs that considerably influence animal productivity as well as the quality of meat and milk.

## 1 Introduction

Forage grasses play a fundamental role in the global beef production chain. Brazil is the country with the greatest emphasis on this sector, being the main exporter of beef and having the largest commercial herd of beef cattle in the world, with approximately 215 million heads distributed in 162 million hectares of pasture (ABIEC, 2019). The main factor that led to this scenario was the beginning of tropical forage breeding in the 1980s in Brazil, which although recent, permitted the country to become the world’s largest exporter of tropical forage seeds (ITC, 2018).

The African forage grass species *Megathyrsus maximus* (Jacq.) B. K. Simon & S. W. L. Jacobs (syn. *Panicum maximum* Jacq.), also known as guinea grass, is one of the most productive forage grasses reproduced by seeds in the Brazilian market and is also grown in other Latin American countries (Jank et al., 2011). It has been used mainly in intensive systems with high-fertility soils (Valle et al., 2009). Moreover, this forage has a high biomass potential and is promising as a biofuel feedstock (Odorico et al., 2018). The polyploidy and domestication process of this forage grass ensure high genetic variability to be explored (Jank et al., 2011); however, a lack of knowledge of the biology and genetics of the species, including its autotetraploidy and facultative apomictic mode of reproduction (Warmke, 1954), may make breeding more difficult and thus stimulate a need to invest in genetic studies.

The polysomic inheritance in autopolyploids makes genetic research difficult because several types of segregation may be involved (Field et al., 2017) and complexity increases as ploidy increases. Thus, in a segregating population, genetic complexity may influence the segregation and the frequencies of expected genotypes (Field et al., 2017), even causing distortions such as double reduction (DR), which is a type of meiosis in which the sister chromatids are duplicated, forming an unexpected combination of gametes. For example, an autotetraploid genotype with the ‘abcd’ alleles at a locus that can form six expected combinations of gametes, namely, ‘ab’, ‘ac’, ‘ad’, ‘bc’, ‘bd’, and ‘cd’, but a homozygous gamete can be generated, e.g., ‘aa’, ‘bb’, ‘cc’, or ‘dd’ (Haldane, 1930; Mather, 1935; Haynes and Douches, 1993). How to accommodate DR and its implications in a breeding program as well as the use of these marker loci in linkage mapping has been discussed for a long time (Butruille and Boiteux, 2000; Luo et al., 2000; Xu et al., 2013; Layman and Busch, 2018; Bourke et al., 2019) for some economically important species, such as potato (Bradshaw, 2007; Bourke et al., 2015) and alfalfa (Julier et al., 2003), but has not yet been reported in guinea grass.

Linkage maps have been used as the primary source of genetic information for nonmodel species that do not have their genome sequenced, such as *M. maximus*. The construction of dense linkage maps allows the identification of the structure and evolution of the genome by mapping traits with polygenic and monogenic inheritance and may even contribute to the assembly of the genome of a species (Doerge, 2002; Flint-Garcia et al., 2003; Luo et al., 2004). The majority of linkage maps available for autotetraploids are based on single-dose segregating markers for a parent (Aaaa x aaaa) or a single dose for both parents (Aaaa x Aaaa). Despite the use of complex statistical methods to obtain integrated maps that combine information from both marker patterns, these maps cover only part of the genome because higher-dose markers (Aaaa, AAAa, and AAAA) are not included. This limitation results in a considerable loss of genetic information. To overcome this limitation, a new approach allows the assignment of allele dosage information for single nucleotide polymorphism (SNP) markers through exact allele sequencing depth, which generates linkage maps from markers in multiple doses with higher quality, more information and greater applicability, including more efficient detection of loci related to traits of economic importance (Serang et al., 2012; Hackett et al., 2014; Pereira et al., 2018).

Mapping of the apomixis region is extremely important for the genetic breeding of *M. maximus* and other forage grasses, such as *Urochloa* spp., *Paspalum* spp., and *Cenchrus ciliaris*. These species undergo asexual propagation by seeds (Nogler, 1984), which allows the fixation of hybrid vigor in apomictic individuals and their use in the creation of uniform pastures (Jank et al., 2011). Experimental field data showed that apomixis in tropical forage grasses follows 1:1 Mendelian segregation, indicating monogenic inheritance (Savidan, 1981; Valle et al., 1994; Chen et al., 2000; Savidan, 2000), although recent evidence suggests that this reproductive mode should be treated as a quantitative trait (Marcón et al., 2019). Molecularly, the apospory-specific genomic region (ASGR), which is responsible for apospory, is highly conserved among apomictic species (Gualtieri et al., 2006). The influence of some factors, such as epigenetics (Kumar, 2017), the presence of retrotransposons (Akiyama et al., 2011), and gene duplication (Conner et al., 2008), in this region has been reported for other species. Due to the laborious and time-consuming methods required to phenotype apomixis, several studies on forage grasses have searched for markers intrinsically linked to the chromosomal region for this trait (Pessino et al., 1998; Ebina et al., 2005; Bluma-Marques et al., 2014; Thaikua et al., 2016; Vigna et al., 2016; Worthington et al., 2016); however, a 100% efficient marker for use in *M. maximus* breeding programs has not yet been identified.

In guinea grass there are no mapping studies of loci related to complex traits, such as those involved in forage yield and nutritional quality. Forage yield results from the continuous emission of leaves and tillers, ensuring the restoration of the leaf area after grazing in perennial pastures. Additionally, the nutritional value of a forage is directly related to animal performance and is measured by the crude protein, in vitro digestibility, neutral and acid detergent fiber, and lignin percentages (Jank et al., 2011). Thus, the mapping of these and other important quantitative trait loci (QTLs) may provide information about the genetic architecture of traits and assist in new strategies for breeding programs of *M. maximus*.

In this context, given the importance of new genomic studies in guinea grass for both biological knowledge and support for breeding programs, our goals were to (i) construct an integrated consensus linkage map from a full-sib progeny of *M. maximus* using allele dosage information, (ii) detect QTLs related to important agronomic and nutritional traits in this progeny, (iii) map the apo-locus, and (iv) search for similarity in regions of the markers adjacent to the QTLs and apomixis locus in *Arabidopsis thaliana, Panicum virgatum* and *Oryza sativa*.

## 2 Material and methods

### 2.1 Plant material

A full-sib progeny of 136 F_1_ hybrids was obtained from a cross between an apomictic genotype of *M. maximus* cv. Mombaça and a sexual genotype, S10. The sexual genotype was derived from sexual x apomictic crosses of an original diploid sexual plant that was duplicated with colchicine (Savidan and Pernès, 1982); thus, both parents were autotetraploid (2n = 4x = 32) (Savidan et al., 1989). In addition to the reproductive mode, the parents have contrasting agronomic and nutritional quality traits (Braz et al., 2017). DNA extraction followed the protocol described by Doyle and Doyle (1987), with modifications. DNA samples were visualized on 2% agarose gels to check their quality and integrity, and their concentrations were estimated using a Qubit 3.0 fluorometer (Thermo Scientific, Wilmington, USA).

### 2.2 Experimental design

A field experiment following an augmented block design (ABD) with 160 regular treatments (full-sib progeny) and two checks (the parents ‘Mombaça’ and S10) distributed in eight blocks with two whole replicates was performed at Embrapa Beef Cattle (Brazilian Agricultural Research Corporation), in Campo Grande city, Mato Grosso do Sul state, Brazil (20°27’S, 54°37’W, 530 m). Each block consisted of a total of 22 plots (20 individuals and two checks).

Each plant was evaluated for agronomic and nutritional quality traits, totaling 22 traits: i) agronomic traits: green matter (GM - g/plant), total dry matter (TDM - g/plant), leaf dry matter (LDM - g/plant), stem dry matter (SDM - g/plant), regrowth capacity (RC), and percentage of leaf blade (PLB - %) and ii) nutritional quality traits for the leaf and stem: organic matter (OM_L and OM_S, respectively - %), crude protein (CP_L and CP_S - %), in vitro digestibility of organic matter (IVD_L and IVD_S - %), neutral detergent fiber (NDF_L and NDF_S - %), acid detergent fiber (ADF_L and ADF_S - %), cellulose (CEL_L and CEL_S - %), silica (SIL_L and SIL_S - %), and permanganate lignin (PL_L and PL_S - %). The agronomic traits were evaluated for six harvests (three harvests in 2013 and three harvests in 2014), but RC was evaluated for only three harvests (one harvest in 2013 and two harvests in 2014). The nutritional quality traits were evaluated for only one harvest in 2014.

### 2.3 Statistical analysis of phenotypic data

Descriptive analyses were performed, and the Box-Cox transformation (Box and Cox, 1964) was applied to correct for nonnormality of the residuals. For traits with multiple harvests (agronomic traits), we fitted the following longitudinal linear mixed model:

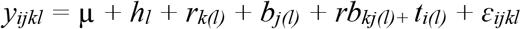

where *y_ijkl_* was the phenotypic value of the *i^th^* treatment in the *j^th^* block and *k^th^* replicate at the *l^th^* harvest; μ was the fixed overall mean; h_l_ was the fixed effect of the *l^th^* harvest (*l* = 1, …, *L*, with *L* = 3 for RC and *L* = 6 for the other traits); *r_k(l)_* was the fixed effect of the *k^th^* replicate (*k* = 1, …, *K*, with *K* = 2) at harvest *l*; *b_j(l)_* was the random effect of the *j^th^* block (*j* = 1, …, *J*, with *J* = 8) at harvest *l*, with 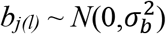; *rb_kj(l)_* was the random interaction effect of replication *k* and block *j* at harvest *l*, with 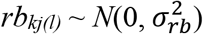; *t_i(l)_* was the effect of the *i^th^* treatment (*i* = 1, …, *I*, with *I* = 162) at harvest *l*; and *ε_ijkl_* was the random environmental error. The treatment effects (*t_i(l)_*) were separated into two groups: *g_i(l)_* was the random effect of the *i^th^* individual genotype (*i* = 1, …, *I_g_*, with *I_g_* = 160) at harvest *l*, and *c_i(l)_* was the fixed effect of the *i^th^* check (*i* = 1, …, *I_c_*, with *I_c_* = 2) at harvest *l*. For genotype effects, the vector ***g*** = (*g_11_*, …, *g_IgL_*)′ was assumed to follow a multivariate normal distribution with a mean of zero and genetic variance-covariance (VCOV) matrix ***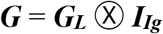***, i.e., ***g*** ~ *N*(**0**, ***G***). For residual effects, the vector ***ε*** = (*ε*_1111_, …, *ε_IJKL_*)′ followed a multivariate normal distribution with a mean of zero and residual VCOV matrix 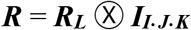, i.e., ***ε*** ~ *MVN*(**0**, ***R***).

The VCOV matrices ***G***_***L***_ and ***R***_***L***_ were analyzed considering seven different structures: identity (ID), diagonal (DIAG), compound symmetry (CS), heterogeneous compound symmetry (CS_Het_), first-order autoregressive (AR1), heterogeneous first-order autoregressive (AR1_Het_), and unstructured (US). First, the genetic VCOV matrix (***G_L_***) was analyzed considering the ID for the residual matrix (***R_L_***), and posteriorly, the residual matrix (***R_L_***) was analyzed considering the selected VCOV matrix for genetic effects. Model selection was performed based on the Akaike information criterion (AIC) (Akaike, 1974) and Schwarz information criterion (SIC) (Schwarz, 1978).

For the nutritional quality traits, we fitted the following linear mixed model:

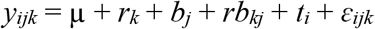

where *y_ijk_* was the phenotypic value of the *i^th^* treatment in the *j^th^* block and *k^th^* replication; μ, *r_k_*, *b_j_*, *rb_kj_*, *t_i_*, and *ε_ijk_* were as described above but not nested within harvest and with 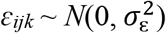. The treatment effects (*t_i_*) were separated into two groups: *g_i_* as a random effect, 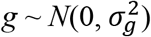, and *c_i_* as a fixed effect. All analyses were performed with the R package ASReml-R (Butler et al., 2009).

The heritability of each trait was calculated using the same model as previously mentioned but considering the ***G_L_*** and ***R_L_*** matrices as the ID. The equation was

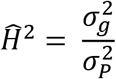

where 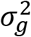 was the genetic variance and 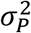 was the phenotypic variance. The network analysis was carried out using the R package ‘qgraph’ (Epskamp et al. 2012).

### 2.4 Identification of the reproductive mode

The apomictic or sexual reproductive mode was determined for 106 hybrids of the progeny (Supplementary Table 1). From the flowers collected during anthesis, we performed an analysis of 30 embryo sacs per hybrid, using the clarified ovary method described by Young et al. (1979). Nomarski differential interference contrast microscopy was used to view the ovaries. A chi-square test was performed to verify the Mendelian segregation of this trait according to the expected model of monogenic inheritance in the base package of R (version 3.5.0) (R Core Team, 2018).

### 2.5 GBS library preparation and sequencing

From the extracted DNA, genotyping-by-sequencing (GBS) libraries were built according to Poland et al. (2012), containing 12 replicates for each parent. A total of 200 ng of genomic DNA per sample was digested with a combination of a rare-cutting enzyme (*Pst*I) and a frequently cutting enzyme (*Msp*I). DNA fragments were ligated to the common and barcode adapters, and the libraries were sequenced as 150-bp single-end reads using the High Output v2 Kit (Illumina, San Diego, USA) for the NextSeq 500 platform (Illumina, San Diego, USA).

### 2.6 SNP calling and allele dosage analysis

First, raw data were checked for quality using NGS QC Toolkit (Patel and Jain, 2012). SNP calling analysis was performed using the TASSEL-GBS v.4 pipeline (Glaubitz et al., 2014) modified for polyploids (Pereira et al., 2018) that use exact read depths. Default parameters changed for the minimum number of times a GBS tag must be present to 5 and the minimum count of reads for a GBS tag to 2. This pipeline requires a genome as a reference for SNP calling, but no genome sequence of *M. maximus* is available. To overcome this limitation, the switchgrass genome (*P. virgatum* v1.0, produced by the US Department of Energy Joint Genome Institute) available in the Phytozome database (http://phytozome.jgi.doe.gov/) (Goodstein et al., 2012) was chosen because this species is phylogenetically closely related to *M. maximus* (Burke et al., 2016). GBS tags were aligned to the reference genome with Bowtie2 2.3.1 (Langmead et al., 2009) using the following settings: very-sensitive-local, a limit of 20 dynamic programming problems (D) and a maximum of 4 times to align a read (R). Subsequently, only tags that aligned exactly one time were processed. Then, SNP calling was performed under the conditions that the minor allele frequency was greater than 0.05 and the minor allele count was greater than 1,000. Mismatches of duplicated SNPs greater than 0.2 were not merged. Then, in R software (version 3.5.0) (R Core Team, 2018), we selected only the SNPs with a minimum average allele depth equal to or greater than 60 reads. The updog package (Gerard et al., 2018) was used to estimate the allele dosage of these markers, with a fixed ploidy parameter of 4 and the flexdog function considering the F_1_ population model. SNPs with less than 0.15 of the posterior proportion of individuals incorrectly genotyped were selected. GBS sequences of each individual were deposited in the NCBI database under number PRJNA563938.

### 2.7 Quality filtering of SNPs

We removed markers with more than 25% missing data and monomorphic markers manually in R software (version 3.5.0) (R Core Team, 2018). Subsequently, we followed Bourke et al. (2018a) to ensure the retention of reliable markers. We first verified the shifted markers for the polysomic inheritance model from which SNP markers that did not correspond to an expected segregation type were removed. A threshold of 5% was used for missing values per marker and per individual. Duplicated markers, which provided no extra information about a locus, were also removed in this step. Finally, a principal component analysis (PCA) was performed to identify individuals that deviated from the progeny as well as possible clones.

### 2.8 Linkage map construction

A linkage map was construed using TetraploidSNPMap version 3.0 (Hackett et al., 2017), which allows the use of SNP markers with allele dosage data for autotetraploid species. SNP markers were checked with a chi-square test for goodness of fit, and only markers with a simplex configuration value greater than 0.001 and a segregation value greater than 0.01 were selected for mapping. Some unselected markers were classified as having segregation distortion (SD), being incompatible with the parental dosages (NP) and having DR. To order the selected markers, two-point analysis and multidimensional scaling analysis (MDS) were used to calculate recombination fractions and logarithm of odds (LOD) scores. Outlier markers were removed in this step. Some phases of the linked SNPs were inferred by TetraploidSNPMap software, and other phases were determined manually. The integrated consensus map represented by homology groups (HGs) was plotted using MapChart 2.32 (Voorrips, 2002), in which SNP configurations were identified with different colors.

### 2.9 Monogenic and polygenic trait analysis

QTL mapping of six agronomic traits and sixteen nutritional quality traits was performed with TetraploidSNPMap, applying an interval mapping model (Hackett et al., 2014, 2017). Analyses were conducted for each HG separately using three data files: phenotypic trait data, genotypic data and map data with phase information. The phenotypic data for the reproductive mode, i.e., apomictic or sexual, were considered qualitative due to the evaluation method applied; i.e., apomictic individuals were coded as one, and sexual individuals, as zero. The other phenotypic traits were analyzed as quantitative. QTL positions and significance were evaluated with a 1,000 permutation test. A QTL was declared significant if its LOD score was above the 90% threshold. Simple models were tested for each significant QTL to verify the best QTL model. The lowest SIC (Schwarz, 1978) was the criterion used to define the best model. Using TetraploidSNPMap software, if two or more significant QTLs were identified on the same chromosome, only the one with the greatest effect was considered.

### 2.10 Search for similarity in apomixis and QTL regions

We performed a search for similarity of candidate genes located close to the detected QTL/apomixis locus regions. Using the switchgrass genome as a reference and based on chromosomal locations of the markers adjacent to the detected QTLs and apomixis locus, we aligned the sequences found in 100-kb regions with Basic Local Alignment Search Tool (BLAST) (e-value cutoff of 1e-0.5) against the *A. thaliana* and *O. sativa* genomes through the JBrowse tool in Phytozome (http://phytozome.jgi.doe.gov/) (Goodstein et al., 2012).

## 3 Results

### 3.1 Phenotypic data

Different VCOV matrices were selected for the agronomic and nutritional traits (Supplementary Tables 2 and 3). When the AIC and SIC were not in agreement, we selected the matrices based on the largest difference between the models. For example, considering the GM trait and the ***G_L_*** matrix, US was selected based on the AIC, and CS was selected based on the SIC (US had 567.32 for the AIC and 702.21 for the SIC, and CS had 581.66 for the AIC and 598.53 for the SIC). The differences between these two selected models were 14.34 for the AIC (581.66-567.32) and 103.68 for the SIC (702.21-598.53). As the SIC produced the largest difference, this criterion was used, and the CS matrix was selected for the ***G_L_*** matrix of the GM trait. The heritabilities ranged from 0.19 (PLB) to 0.64 (GM) for the agronomic traits and from 0.06 (SIL_S) to 0.31 (OM_L and CEL_S) for the nutritional quality traits (Table 1). Box-Cox transformation was performed for GM, TDM, LDM, SDM, RC, PLB, OM_L, OM_S, CP_S, IVD_L, IVD_S, NDF_S, ADF_L, CEL_L, SIL_L, SIL_S, PL_L, and PL_S. The correlations between the agronomic and nutritional traits are presented in Figure 1.

**Table 1.**
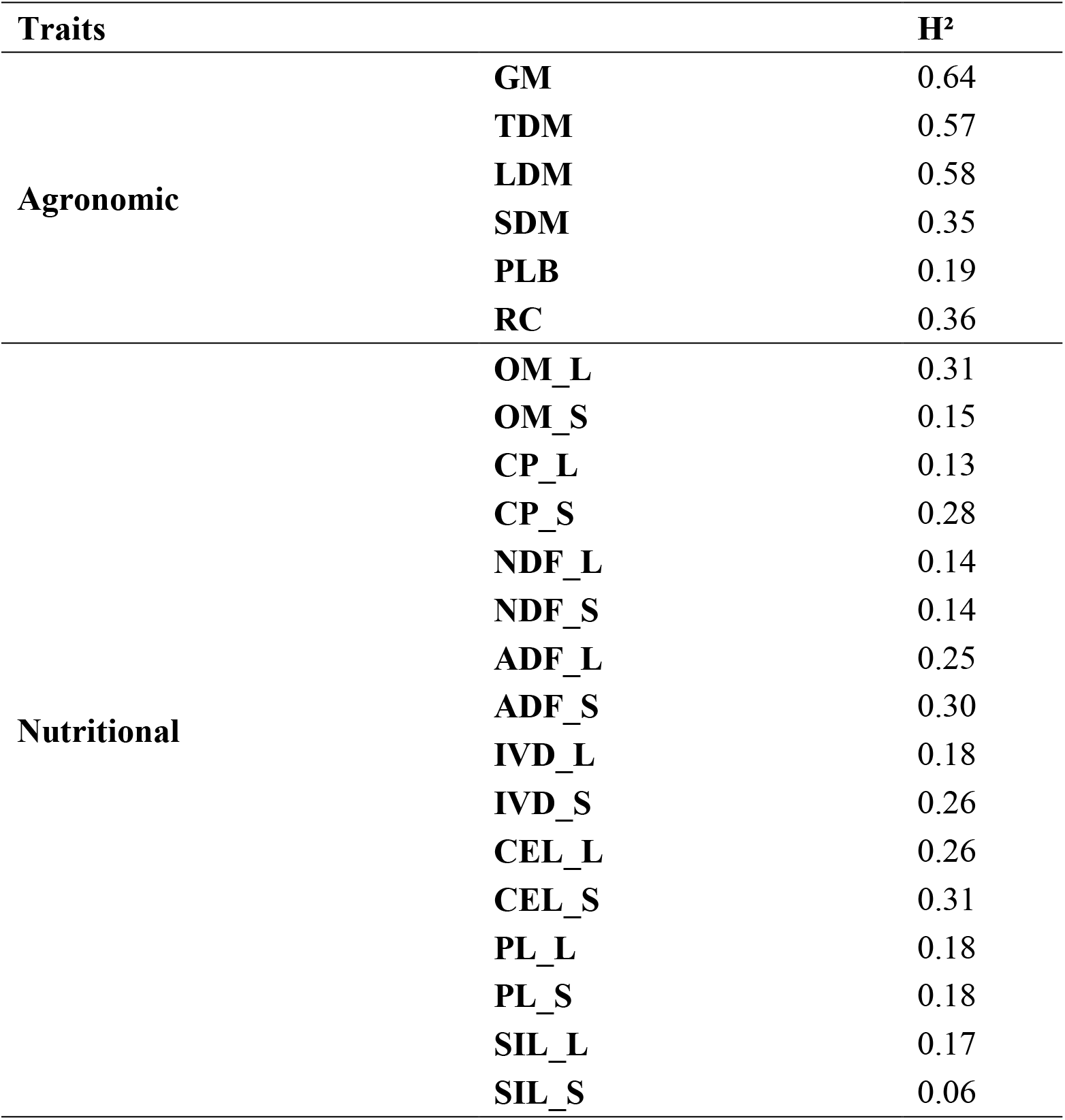
Broad-sense heritability obtained for the agronomic and nutritional traits for a mapping population of guinea grass (*Megathyrsus maximus*).

**Figure 1.**
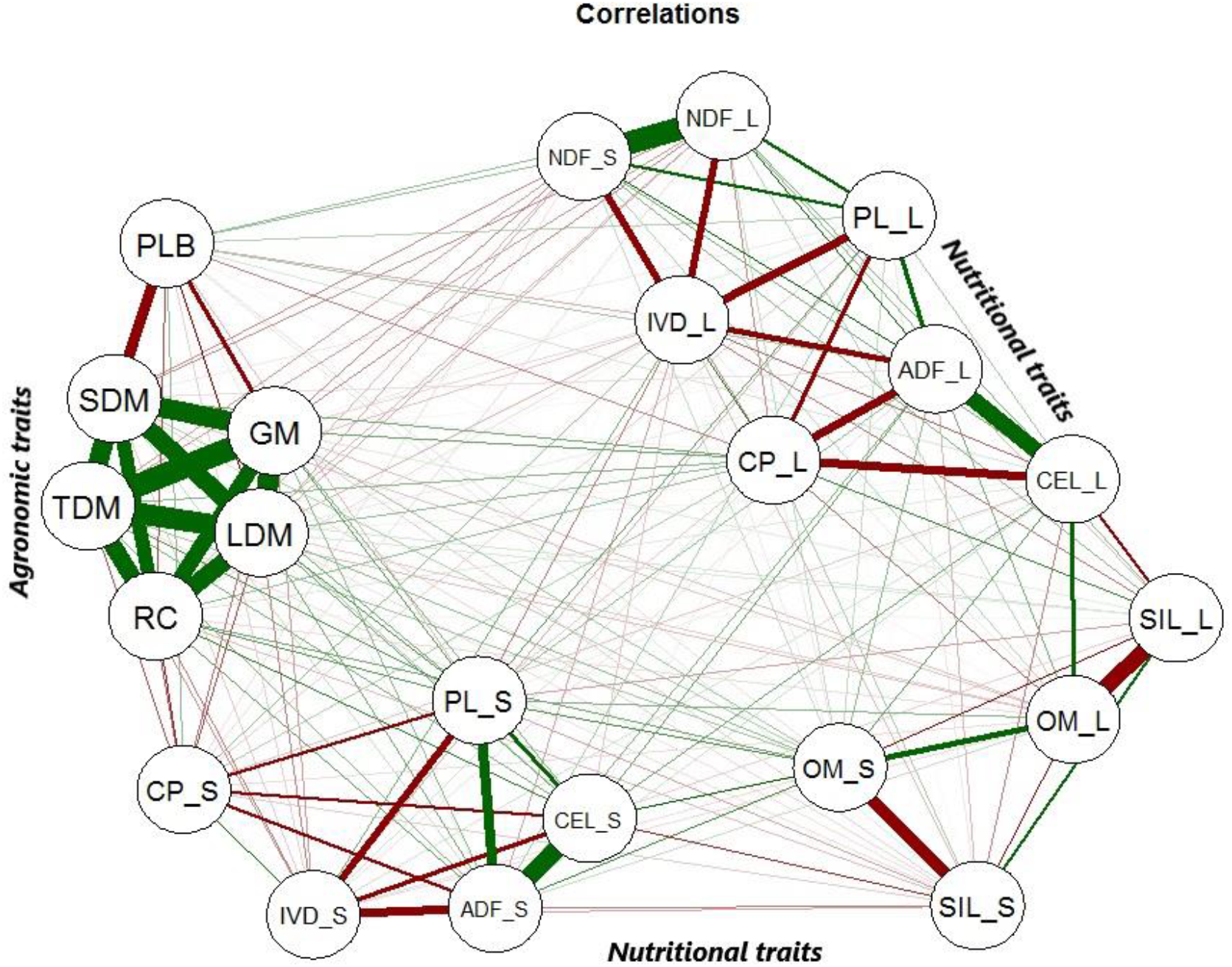
Correlations among all phenotypic traits from guinea grass mapping population. The green lines correspond to positive correlations and the red lines correspond to negative correlations between the traits.

### 3.2 Genotyping and linkage analysis

A total of 23,619 SNPs were identified with the alignment using the *P. virgatum* genome as the reference. After the genotypic analysis, 2,804 SNPs with an average allele depth greater than 60 were selected for further filtering. After allele dosage estimation, 275 SNPs were discarded, in addition to 15 monomorphic markers and four markers with missing data identified manually in R software. Using the updog package, we obtained the allele dosage information for 2,510 SNPs, considering an autotetraploid species (Table 2). Simplex (AAAA × AAAB, ABBB × BBBB) was the most commonly found configuration, with 1,654 markers, followed by the duplex (AAAA × AABB, AABB × BBBB) and double-simplex (AAAB × AAAB, ABBB × ABBB) configurations, with 334 and 226 SNPs, respectively. The simplex-duplex (AAAB × AABB) configuration was the most represented among the higher dosages, with 155 markers, and the duplex-simplex (AABB × ABBB) configuration was the least represented, with only six SNPs. Analysis with the polymapR package revealed that all offspring were 95% compatible with the parents. Two individuals (B126 and B127) were very genetically similar to the parent S10 and were removed, and one clone (C49) of individual C44 was also removed (Figure 2). After this, we discarded another 37 SNPs with missing values and 1,151 duplicated SNPs. This filtering resulted in 132 genotyped offspring with 1,322 markers that were used at the beginning of the linkage analysis in TetraploidSNPMap software. In this step, incompatible markers with the parental allele dosages, SNPs with DR and markers showing SD were not considered. In addition, two-point analysis identified 149 SNPs as duplicated, and MDS analysis identified 27 outliers, which were then excluded. In total, 858 reliable SNPs were included in the linkage map. The apomictic parent, cv. Mombaça, presented 368 exclusive alleles; the sexual parent, S10, presented 275 exclusive alleles; and the two parents shared 215 alleles. TetraploidSNPMap software was used again to rank the 2,510 SNPs based on their expected segregation. The analysis resulted in a total of 114 SNPs with SD, 183 with NP and 243 with DR (Table 2).

**Table 2.** Distribution of SNP markers among genotype classes for a mapping population of guinea grass (*Megathyrsus maximus*).

| Marker class | Genotype of the parents | Segregation ratio | Number | DR | NP | Distorted | Mapped |
| --- | --- | --- | --- | --- | --- | --- | --- |
| **Null** | AAAA x BBBB | 0 | 0 | 0 | 0 | 0 | 0 |
| **Simplex** | AAAA x AAAB, ABBB x BBBB | 1:1 | 1,654 | 111 | 125 | 18 | 491 |
| **Duplex** | AAAA x AABB, AABB x BBBB | 1:4:1 | 334 | 0 | 58 | 40 | 136 |
| **Triplex** | AAAA x ABBB, AAAB x BBBB | 1:1 | 35 | 18 | 0 | 0 | 17 |
| **Double-Simplex** | AAAB x AAAB, ABBB x ABBB | 1:2:1 | 226 | 60 | 0 | 5 | 106 |
| **X-Double-Simplex** | AAAB x ABBB | 1:2:1 | 26 | 16 | 0 | 0 | 9 |
| **Simplex-Duplex** | AAAB x AABB | 1:5:5:1 | 155 | 38 | 0 | 9 | 72 |
| **Duplex-Simplex** | AABB x ABBB | 1:5:5:1 | 6 | 0 | 0 | 0 | 1 |
| **Double-Duplex** | AABB x AABB | 1:8:18:8:1 | 74 | 0 | 0 | 42 | 26 |
| **Total** |  |  | **2510** | **243** | **183** | **114** | **858** |

Significant distortion at ( $P < 0.001$ ) and ( $P < 0.01$ ) for simplex and higher-dosage markers. DR corresponds to double reduction and NP are incompatible SNPs with the parental dosages. SNPs with missing data, outliers and duplicates were not considered.

**Figure 2.**
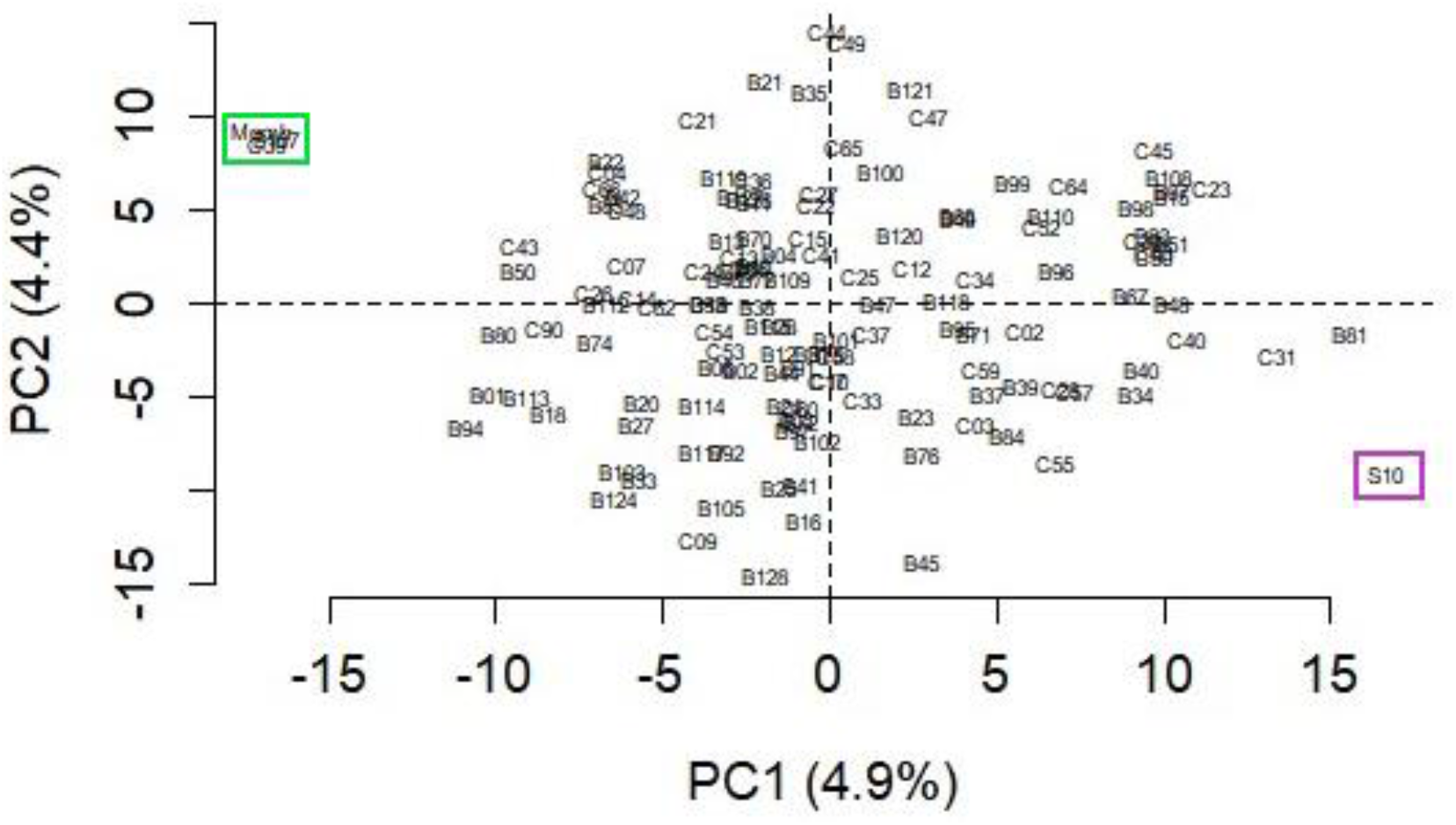
Principal component analysis representing the genetic diversity among the progeny and the parents of the mapping population of guinea grass (*Megathyrsus maximus*). Parents were highlighted in pink (S10) and green (cv. Mombaça).

### 3.3 Linkage map

We constructed an integrated consensus linkage map consisting of 858 SNP markers distributed over 756.69 cM with all possible allele dosage configurations for an autotetraploid species (Table 2, Figure 3 and Supplementary Table 4). Considering all integrated consensus HGs, an average density of ~1.13 SNPs/cM was obtained. The largest HG was VII, with 159 SNPs distributed over 108.573 cM, and the smallest HG was VIII, with 49 SNPs present over 70.05 cM. The interlocus intervals were relatively small, with a minimum value of 0.003 cM for the majority of the HGs (I, II, IV, VI and VII) and a maximum of 8.65 cM and 7.24 cM on HG V and HG I, respectively (Table 3). Among the markers, we identified approximately 30 double-duplex markers (AABB x AABB), which contains all types of doses for autotetraploid progenies (Table 2, Figure 3 and Supplementary Table 4) (Hackett et al., 2014).

**Table 3.** Summary of the linkage map of guinea grass (*Megathyrsus maximus*) obtained with the S10 × cv. Mombaça population.

| HG* | No. mapped SNPs | Length (cM) | Smallest interval (cM) | Longest interval (cM) |
| --- | --- | --- | --- | --- |
| **I** | 144 | 101.74 | 0.015 | 7.24 |
| **II** | 128 | 103.53 | 0.031 | 3.43 |
| **III** | 85 | 94.06 | 0.027 | 5.24 |
| **IV** | 115 | 87.66 | 0.003 | 6.34 |
| **V** | 104 | 96.08 | 0.008 | 8.65 |
| **VI** | 74 | 94.99 | 0.013 | 6.42 |
| **VII** | 159 | 108.57 | 0.009 | 5.63 |
| **VIII** | 49 | 70.05 | 0.072 | 6.93 |
| **Total** | **858** | **756,69** |  |  |

\*Homology group

**Figure 3.**
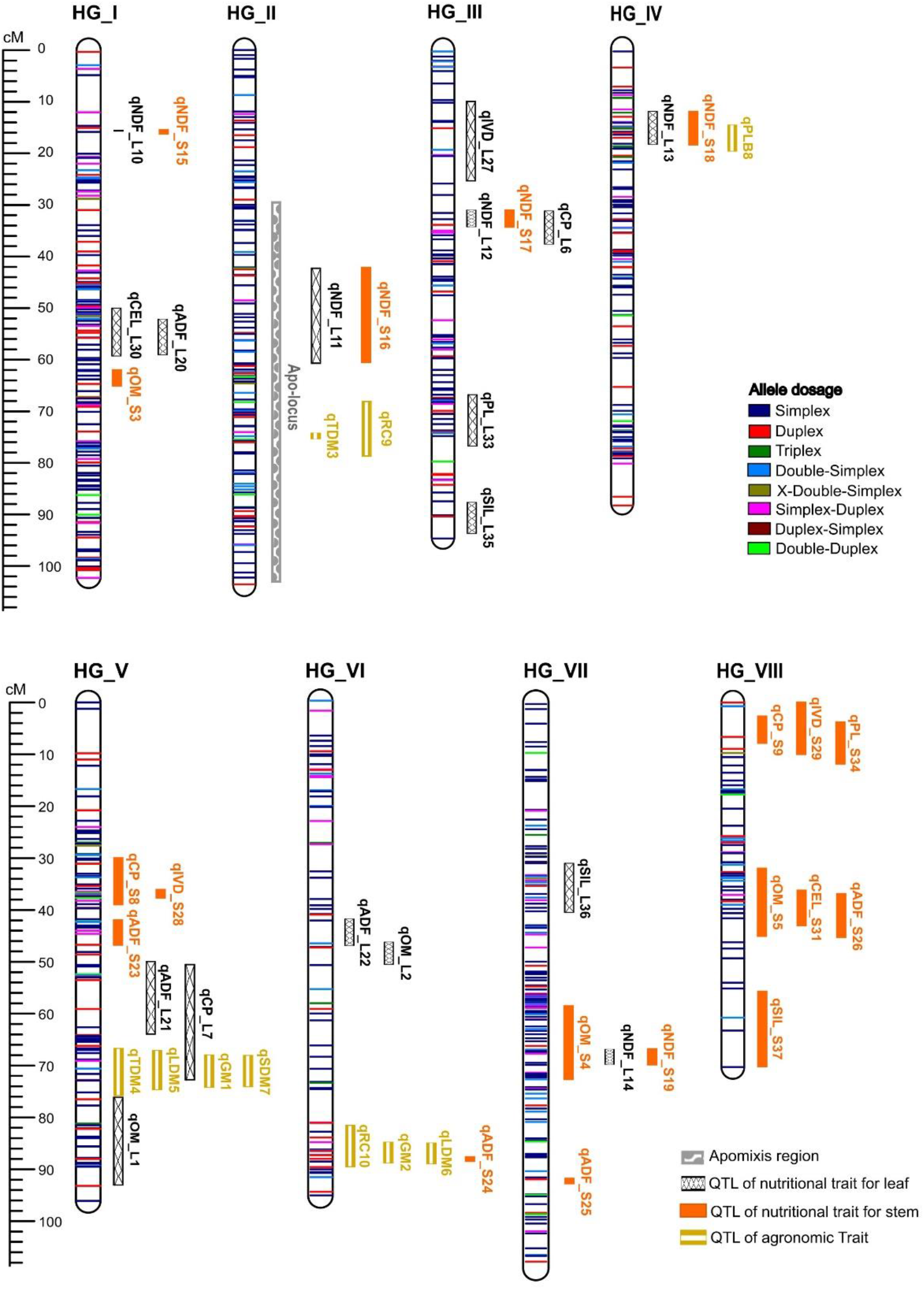
Linkage map constructed for guinea grass (*Megathyrsus maximus*) using SNPs with allele dosage information, including representation of intervals of the highest peaks of QTLs.

### 3.4 Apospory mapping

The mode of reproduction of 106 hybrids in the mapping population was determined and indicated that apospory had a segregation ratio of 1:1 based on a chi-square test (X^2^ = 5.43, p ≥ 0.01), consistent with the model expected for monogenic inheritance. The apo-locus was mapped in the HG II at a peak position of 65 cM, with a high LOD score of 50.06 (Supplementary Figure 1). More than 80% of the phenotypic variation in apomictic reproductive mode was explained, and the simple models that best classified the apo-locus as a simplex genotype (BBBB x ABBB). As expected, this locus was linked to the apomictic parent, cv. Mombaça. Two SNP markers exclusive to this parent, namely, S_14_29023868 and S_10_48091934, were linked to the apo-locus at 0.8 cM (Figure 3).

### 3.5 Agronomic trait mapping

Ten significant QTLs were mapped for GM, TDM, LDM, SDM, PLB and RC, which were distributed in HGs II, IV, V and VI (Table 4 and Supplementary Figure 2). Here, we will discuss the QTLs that were identified for the main traits targeted in the *M. maximus* breeding program. TDM was associated with two QTLs in HGs II (qTDM3) and V (qTDM4) with LOD scores of 3.4 and 4.4 and that explained 5.8 and 9.4% of the phenotypic variation, respectively. For LDM, we detected two QTLs in HGs V (qLDM5) and VI (qLDM6) with LOD scores of 3.9 and 3.0 and that explained 7.9 and 4.5% of the phenotypic variation, respectively. PLB was associated with one QTL in HG IV (qPLB8) with a LOD score of 3.8 and that explained 6.8% of the phenotypic variation. We identified two QTLs for RC in HGs II (qRC9) and VI (qRC10) with LOD scores of 4.7 and 3.8 and that explained 10.4% and 6.3% of the phenotypic variation, respectively.

**Table 4.** QTLs identified for agronomic traits from the sexual genotype S10 and apomictic cv. Mombaça of guinea grass (*Megathyrsus maximus*).

| Agronomic trait | QTL | HG | Position (cM) | LOD | R<sup>2</sup> | Parents |
| --- | --- | --- | --- | --- | --- | --- |
| **GM** | qGM1 | V | 71.0 | 3.56 | 6.78 | both |
|  | qGM2 | VI | 87.0 | 2.91 | 4.29 | Mombaça |
| **TDM** | qTDM3 | II | 75.0 | 3.37 | 5.79 | both |
|  | qTDM4 | V | 70.0 | 4.41 | 9.40 | S10 |
| **LDM** | qLDM5 | V | 70.0 | 3.90 | 7.93 | S10 |
|  | qLDM6 | VI | 88.0 | 2.97 | 4.53 | Mombaça |
| **SDM** | qSDM7 | V | 71.0 | 3.73 | 7.24 | Mombaça |
| **PLB** | qPLB8 | IV | 17.0 | 3.81 | 6.84 | both |
| **RC** | qRC9 | II | 75.0 | 4.73 | 10.37 | S10 |
|  | qRC10 | VI | 87.0 | 3.77 | 6.25 | Mombaça |

The simple models that best represented the QTLs for each agronomic trait were identified based on the lowest SIC values. The QTL with the greatest effect on TDM, which was present in HG VI, followed a duplex model (AAaa x aaaa) with an additive effect of the A allele from the S10 parent. The same model was verified for LDM with qLDM5, a QTL associated with the S10 parent, and qLDM6, a QTL associated with the cv. Mombaça parent. The QTL with the smallest effect for TDM and the single QTL for PLB followed the double-simplex model (Aaaa x Aaaa) with an additive effect of both parents. The two QTLs detected for RC followed the simplex model, with the S10 parent contributing the allele for qRC9 and the ‘Mombaça’ parent contributing the allele for qRC10.

### 3.6 Nutritional trait mapping

Thirty-six significant QTLs were identified for OM, CP, NDF, ADF, IVD, PL, CEL, and SIL for the leaf and stem. These QTLs were distributed in all HGs (Table 5 and Supplementary Figure 3). For CP_L, two QTLs were detected in HGs III (qCP_L6) and V (qCP_L7) with LOD scores of 3.4 and 4.3 and that explained 5.1% and 9.7% of the phenotypic variation, respectively. For CP_S, two QTLs were detected in HGs V (qCP_S8) and VIII (qCP_S9) with LOD scores of 3.8 and 2.8 and that explained 8.4% and 3.2% of the phenotypic variation, respectively. For NDF_L and NDF_S, we detected five QTLs each, all at the same positions in HGs I (qNDF_L10/qNDF_S15), II (qNDF_L11/qNDF_S16), III (qNDF_L12/qNDF_S17), IV (qNDF_L13/qNDF_S18), and VII (qNDF_L14/qNDF_S19). The QTLs with the largest effect on NDF were found in HGs II and IV and were identified with LOD scores of 5.5 and 4.3, respectively; qNDF_L11/qNDF_S16 in HG II explained more phenotypic variation (12%). For the IVD_L trait, only one QTL was found in HG III (qIVD_L27) with a LOD score of 3.8 and that explained 5.9% of the phenotypic variation. For IVD_S, two QTLs were obtained in HGs V (qIVD_S28) and VIII (qIVD_S29) with LOD scores of 3.6 and 3.9 and explaining 7.3 and 5.8% of the phenotypic variation, respectively. We detected a single QTL for PL_L located in HG III (qPL_L33) with a LOD score of 3.9, and it explained 6.2% of the phenotypic variation. Additionally, PL_S was associated with only one QTL, which was identified in HG VIII (qPL_S34) with a LOD score of 2.7 and explained 2.8% of the variation in the trait.

**Table 5.**
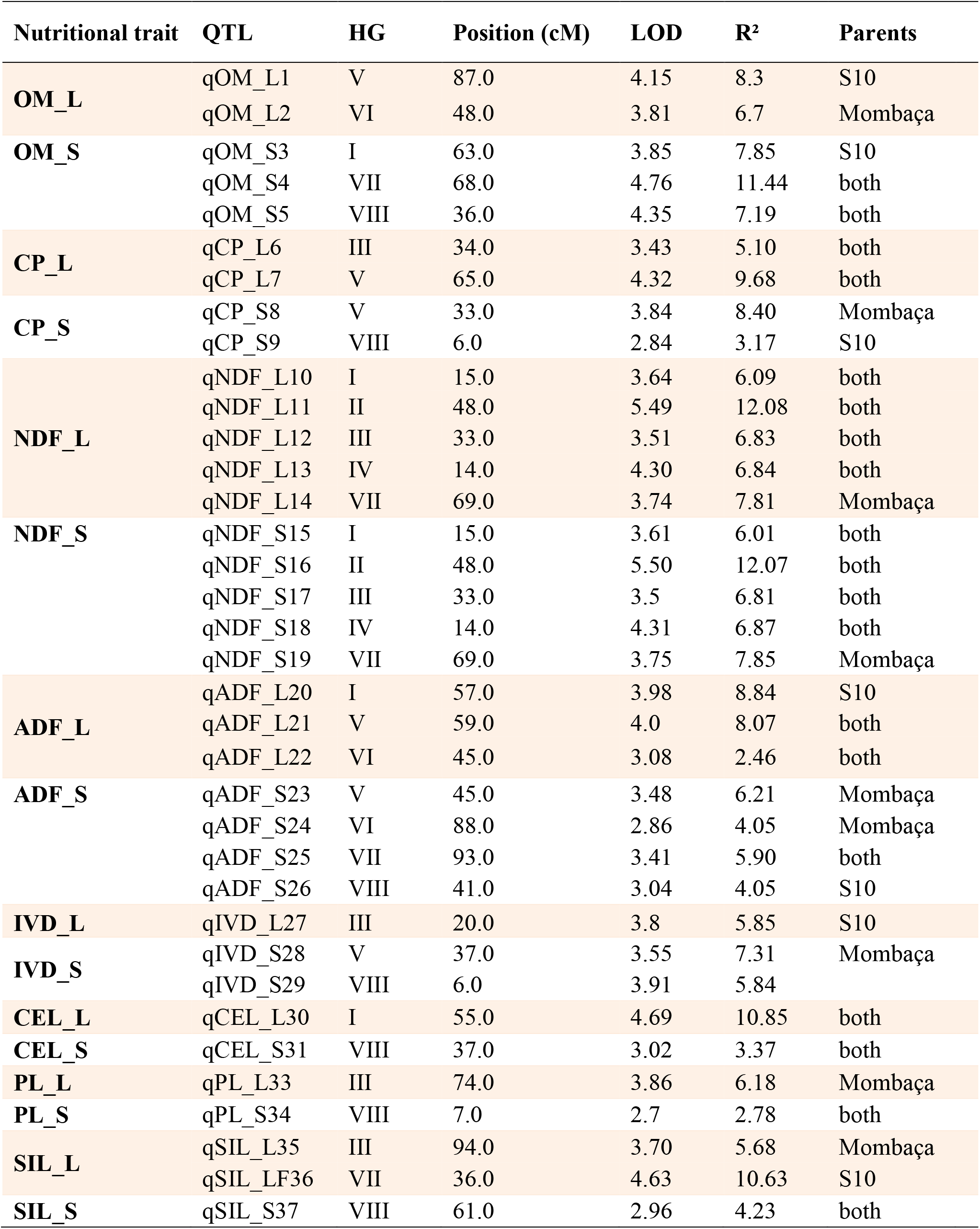
QTLs identified for traits related to nutritional quality from the sexual genotype ‘S10’ and apomictic cv. ‘Mombaça’ of guinea grass (*Megathyrsus maximus*).

| Nutritional trait | QTL | HG | Position (cM) | LOD | R<sup>2</sup> | Parents |
| --- | --- | --- | --- | --- | --- | --- |
| **OM_L** | qOM_L1 | V | 87.0 | 4.15 | 8.3 | S10 |
|  | qOM_L2 | VI | 48.0 | 3.81 | 6.7 | Mombaça |
| **OM_S** | qOM_S3 | I | 63.0 | 3.85 | 7.85 | S10 |
|  | qOM_S4 | VII | 68.0 | 4.76 | 11.44 | both |
|  | qOM_S5 | VIII | 36.0 | 4.35 | 7.19 | both |
| **CP_L** | qCP_L6 | III | 34.0 | 3.43 | 5.10 | both |
|  | qCP_L7 | V | 65.0 | 4.32 | 9.68 | both |
| **CP_S** | qCP_S8 | V | 33.0 | 3.84 | 8.40 | Mombaça |
|  | qCP_S9 | VIII | 6.0 | 2.84 | 3.17 | S10 |
| **NDF_L** | qNDF_L10 | I | 15.0 | 3.64 | 6.09 | both |
|  | qNDF_L11 | II | 48.0 | 5.49 | 12.08 | both |
|  | qNDF_L12 | III | 33.0 | 3.51 | 6.83 | both |
|  | qNDF_L13 | IV | 14.0 | 4.30 | 6.84 | both |
|  | qNDF_L14 | VII | 69.0 | 3.74 | 7.81 | Mombaça |
| **NDF_S** | qNDF_S15 | I | 15.0 | 3.61 | 6.01 | both |
|  | qNDF_S16 | II | 48.0 | 5.50 | 12.07 | both |
|  | qNDF_S17 | III | 33.0 | 3.5 | 6.81 | both |
|  | qNDF_S18 | IV | 14.0 | 4.31 | 6.87 | both |
|  | qNDF_S19 | VII | 69.0 | 3.75 | 7.85 | Mombaça |
| **ADF_L** | qADF_L20 | I | 57.0 | 3.98 | 8.84 | S10 |
|  | qADF_L21 | V | 59.0 | 4.0 | 8.07 | both |
|  | qADF_L22 | VI | 45.0 | 3.08 | 2.46 | both |
| **ADF_S** | qADF_S23 | V | 45.0 | 3.48 | 6.21 | Mombaça |
|  | qADF_S24 | VI | 88.0 | 2.86 | 4.05 | Mombaça |
|  | qADF_S25 | VII | 93.0 | 3.41 | 5.90 | both |
|  | qADF_S26 | VIII | 41.0 | 3.04 | 4.05 | S10 |
| **IVD_L** | qIVD_L27 | III | 20.0 | 3.8 | 5.85 | S10 |
| **IVD_S** | qIVD_S28 | V | 37.0 | 3.55 | 7.31 | Mombaça |
|  | qIVD_S29 | VIII | 6.0 | 3.91 | 5.84 |  |
| **CEL_L** | qCEL_L30 | I | 55.0 | 4.69 | 10.85 | both |
| **CEL_S** | qCEL_S31 | VIII | 37.0 | 3.02 | 3.37 | both |
| **PL_L** | qPL_L33 | III | 74.0 | 3.86 | 6.18 | Mombaça |
| **PL_S** | qPL_S34 | VIII | 7.0 | 2.7 | 2.78 | both |
| **SIL_L** | qSIL_L35 | III | 94.0 | 3.70 | 5.68 | Mombaça |
|  | qSIL_LF36 | VII | 36.0 | 4.63 | 10.63 | S10 |
| **SIL_S** | qSIL_S37 | VIII | 61.0 | 2.96 | 4.23 | both |

According to the simple model analysis, the best model for the two QTLs of CP_L was the double-simplex model (Aaaa x Aaaa) with an additive effect of both parents. In contrast, qCP_S8 was best represented by a duplex model (AAaa x aaaa) with a dominant effect of cv. Mombaça. The CP_S9 QTL was represented by a double-simplex model with an additive effect of the S10 parent. The QTL with the greatest effect on NDF was best represented by the double-simplex model with an additive effect from both parents. For IVD_L, a simplex allele was verified in the sexual parent (Aaaa x aaaa). The same model was also observed for IVD_S and PL_L, with the A allele from the apomictic parent associated with the trait. The single QTL for PL_S was explained by a double-simplex model with a dominant effect from both parents.

### 3.7 Search for similarity in apomixis and QTL regions

Some genes that contain conserved protein domains were found in regions flanking the apo-locus located in HG II, such as the Spc97/Spc98 family of spindle pole body (SBP) components, whose proteins assist in the control of the microtubule network (Lin et al., 2015), and Inner centromere protein (ARK binding region). These two genes play an important role in the segregation of chromosomes during cell division (Kirioukhova et al., 2011; Lin et al., 2015). In addition, a gene possibly involved in the apomictic reproductive mode was similar to somatic embryogenesis receptor-like kinase 1 (SERK1), which is part of a complex associated with the induction of embryo development (Albrecht et al., 2008).

A total of 23 regions were found due to the overlap of QTLs in common regions in the linkage map. In these regions, 55 different gene families from *P. virgatum*, *A. thaliana* and *O. sativa* were identified. Most of these genes may play important roles in cellular metabolism and may be associated with plant growth and development (Table 6). Further details about the locations of the candidate genes in their respective QTL regions are provided in Supplementary Table 5.

**Table 6.**
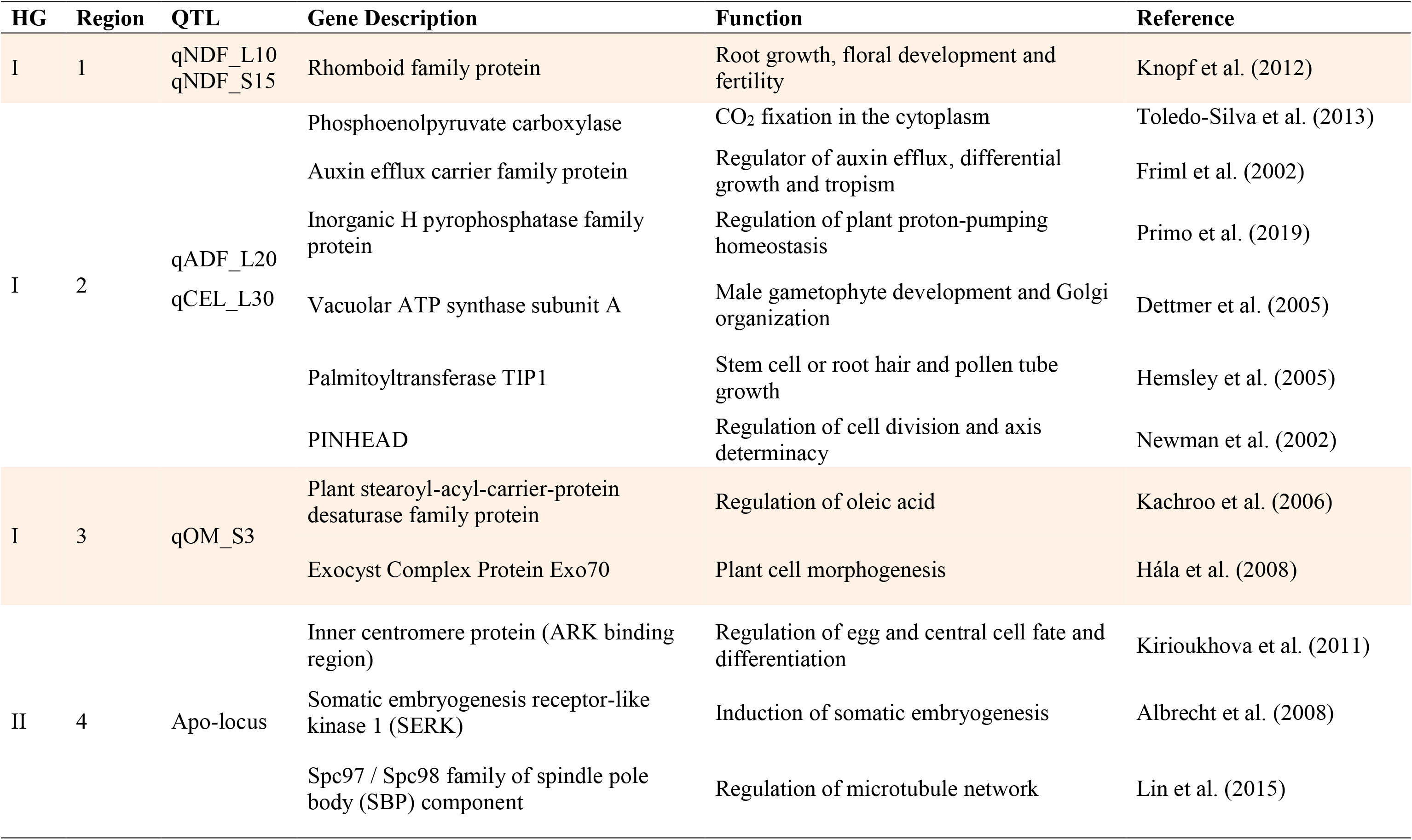
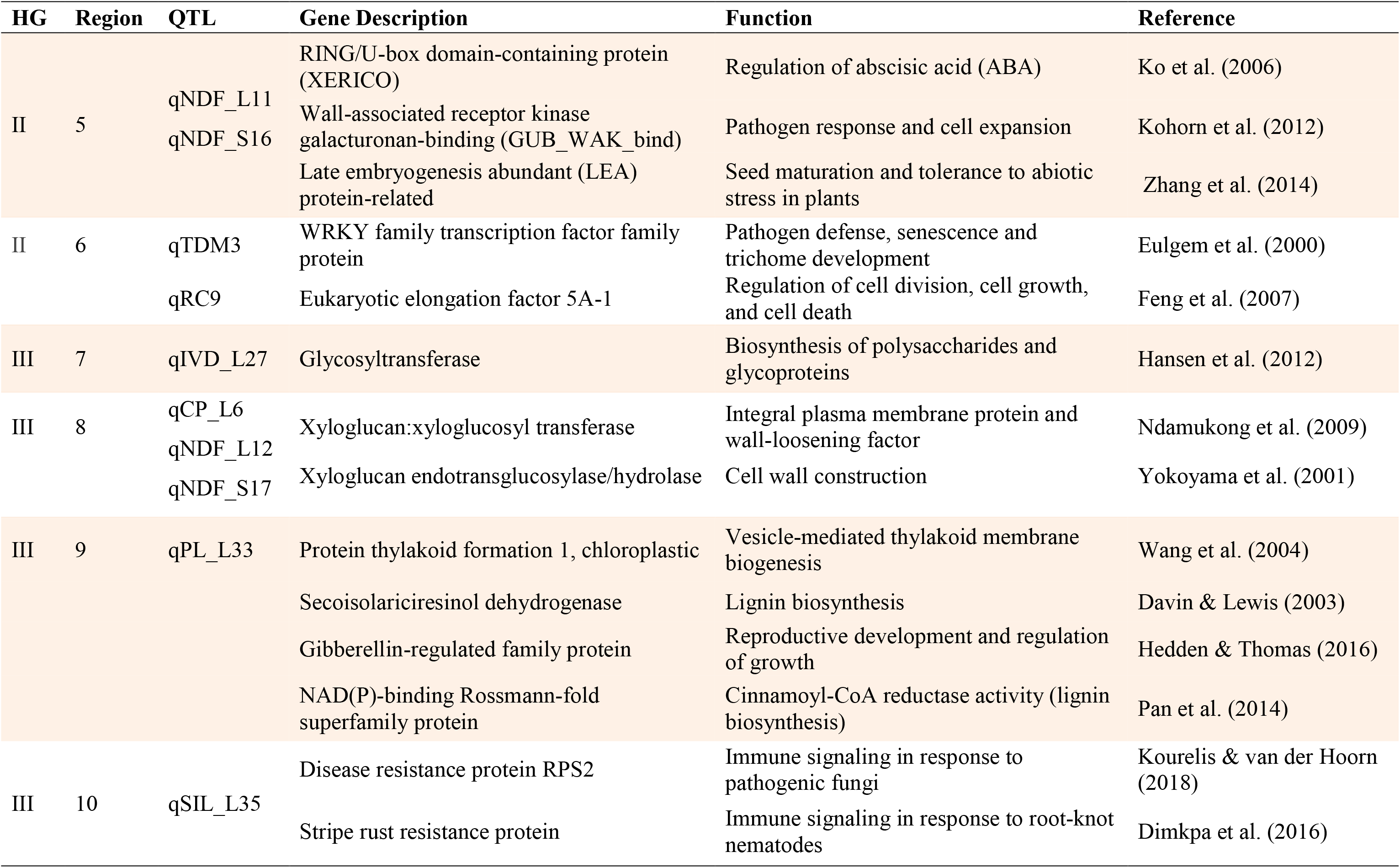
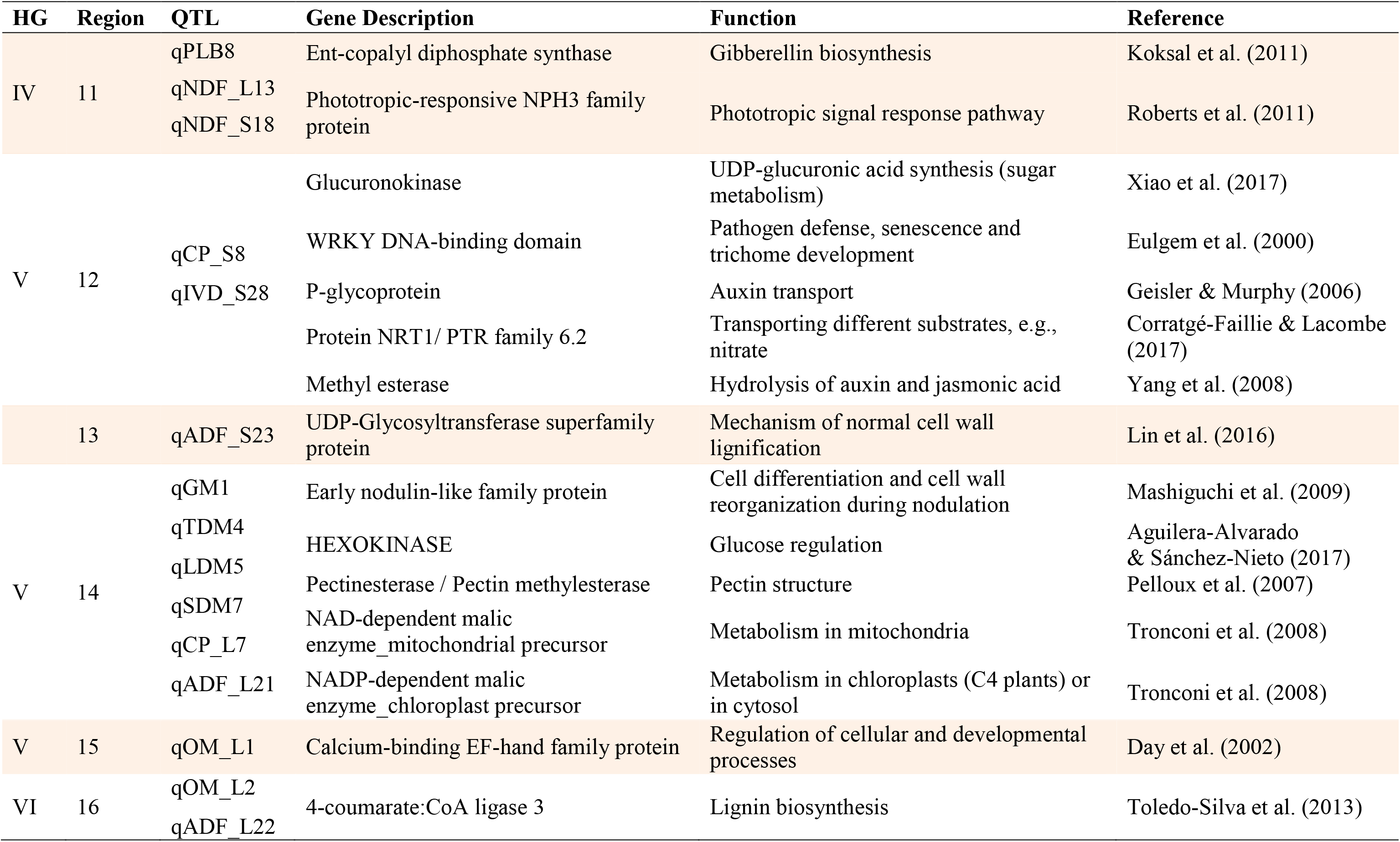
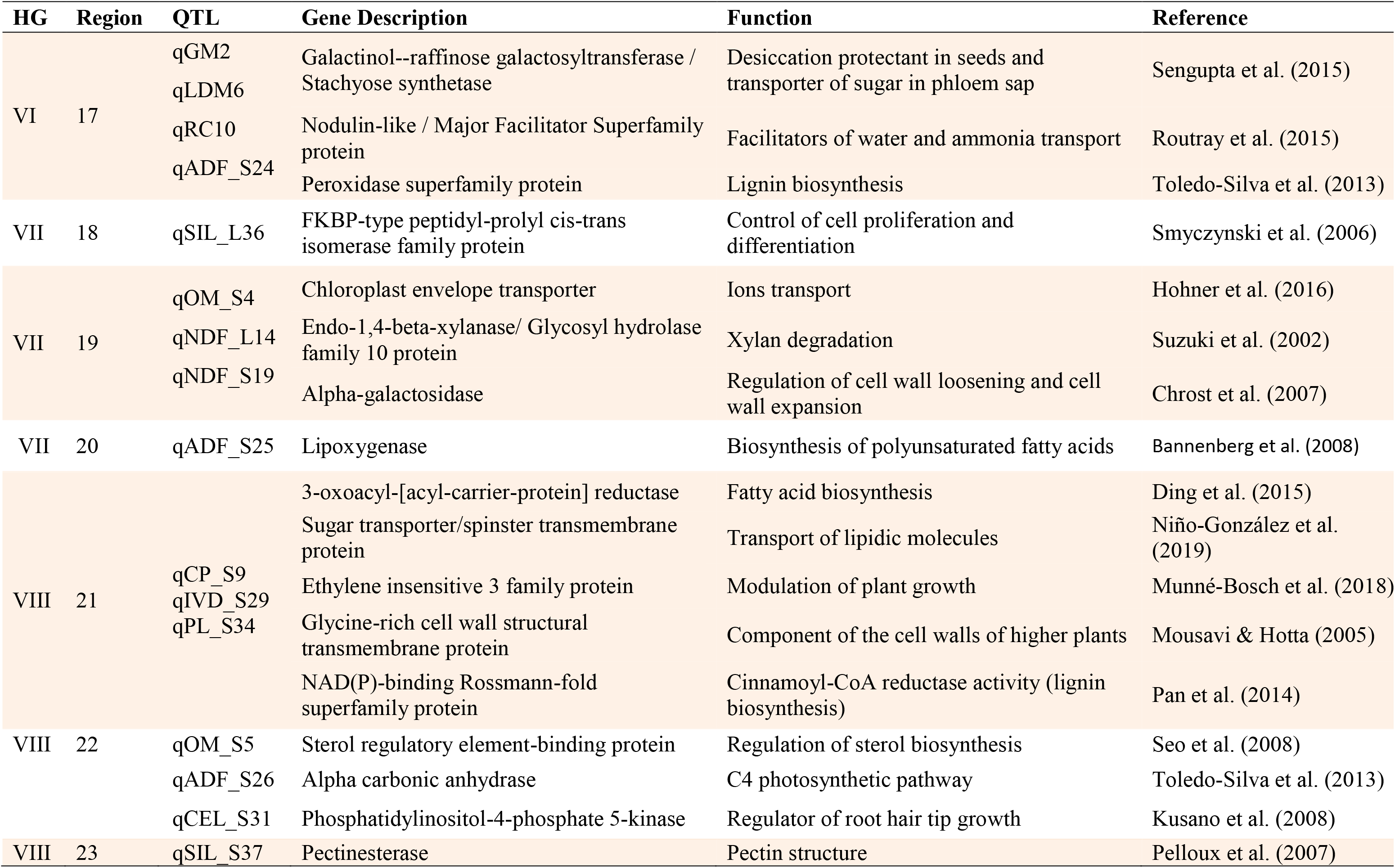
Description and function of the genes identified in apomixis and QTL regions from linkage map of guinea grass.

Among the putative genes, many participate in hormonal signaling pathways, such as Auxin efflux carrier component in HG I - region 2 (qADF_L20/qCEL_L30), which is related to the intercellular directionality of auxin (Friml et al., 2002); RING/U-box (XERICO) in HG II – region 5 (qNDF_L11/ qNDF_S16), which is involved in the regulation of abscisic acid (ABA) (Ko et al., 2006); gibberellin-regulated family members (GAST1, GASR2, GASR3, and GASR9) in HG III – region 9 (qPL_L33), which are associated with the regulation of gibberellic acid (GA) and ABA (Hedden and Thomas, 2016); Ent-copalyl diphosphate synthase in HG IV – region 11 (qNDF_L13/qNDF_S18/qPLB8), which plays a role in GA biosynthesis (Koksal et al., 2011); and Ethylene insensitive 3 (EIN3) in HG VIII – region 21 (qCP_S9/qIVD_S29/ qPL_S34), which is associated with cell growth and senescence processes (Munné-Bosch et al., 2018) (Table 6 and Supplementary Table 5).

In addition, other genes with a role in plant physiology were also identified, such as the gene encoding the enzyme phosphoenolpyruvate carboxylase (PEPC) in HG I – region 2 (qADF_L20/qCEL_L30), whose function is the catalysis of primary metabolic reactions in plants (Toledo-Silva et al., 2013). Glycosyltransferase was found in HG III – region 7 (qIVD_L27) and is associated with the biosynthesis of polysaccharides and glycoproteins (Hansen et al., 2012). Pectinesterase/pectin methylesterase, which was detected in HG V – region 14 (qGM1/qTDM4/qLDM5/qSDM7/qADF_L21/qCP_L7) and HG VIII – region 23 (qSIL_S37) is related to cellular adhesion and stem elongation (Damm et al., 2015). In addition to these genes, we found a chloroplast envelope transporter in HG 7 – region 19 (qOM_S4/qNDF_L14/qNDF_S19). Phosphatidylinositol-4-phosphate 5-kinase, which was present in HG VIII – region 22 (qOM_S5/qADF_S26/qCEL_S31), is involved in coordinating plant growth (Kusano et al., 2008).

Many putative candidate genes present in lignan biosynthesis pathways were verified in some regions of QTLs. Glycosyl hydrolase family 16 was located in HG III - region 8 (qCP_L6/qNDF_L12/qNDF_S17), and the genes associated with secoisolariciresinol dehydrogenase and the NAD(P)-binding Rossmann-fold superfamily were located in region 9 (qPL_L33). UDP-Glycosyltransferase superfamily protein was present in HG V – region 13 (qADF_S23). The 4- coumarate:CoA ligase 3 gene was obtained in HG VI- region 16 (qOM_L2/qADF_L22). The gene encoding NAD(P)-binding Rossmann-fold superfamily was also detected in HG VIII – region 21 (qCP_S9/qIVD_S29/ qPL_S34).

## 4 Discussion

Here, we will discuss the advances resulting from this study, which serves as a model for genetic studies on autotetraploid forage grasses. In addition to the construction of a genetic map for *M. maximus* with the use of allele dosage information, we mapped QTLs for agronomic and nutritional traits, two novel approaches for this species.

### 4.1 Linkage map

We obtained a satisfactory representative linkage map for *M. maximus*, with a large genetic distance observed between the parents, namely, S10 (Figure 2, quadrant I) and cv. Mombaça (Figure 2, quadrant IV). This genetic distance and the distribution of hybrids can be visualized in the PCA performed with the allele dosage information of all individuals (Figure 2). The crossing of contrasting parents allows maximum recombination between loci, which is a fundamental principle for observing the segregation of traits in hybrids and promoting the detection of QTLs (Wang et al., 2017).

Because *M. maximus* does not yet have a sequenced genome, we aligned our GBS reads to the allotetraploid genome of *P. virgatum*, which is a species closely phylogenetically related to *M. maximus* (Burke et al., 2016), since *M. maximus* previously belonged to the *Panicum* genus. This genetic proximity was confirmed based on the consistent distribution of the markers in the genetic map (Figure 3 and Supplementary Table 4). In addition, the use of a tetraploid genome, such as that of *P. virgatum*, may be more informative than the use of diploid genomes, considering our tetraploid mapping population, as there is a greater possibility of similar chromosomal rearrangements between these species (Daverdin et al., 2015). It is common to find a smaller number of SNPs in linkage maps of forage grass species without an assembled genome, as observed for common millet (*Panicum miliaceum*) (Rajput et al., 2016) and signalgrass (*Urochloa decumbens*) (Ferreira et al., 2019), than in those of species with a reference genome, such as foxtail millet (*Setaria italica*) (Jia et al., 2017) and pearl millet (*Pennisetum glaucum*) (Punnuri et al., 2016). Another alternative is the use of “pseudogenomes” assembled from GBS tags to include unique alleles of a species, as in *Paspalum vaginatum*, but the number of markers remained relatively small (Qi et al., 2019). Therefore, the number of SNPs obtained from our nonredundant tags does not preclude the need for a sequenced genome of this species, and our linkage map can contribute to the assembly of this reference genome of guinea grass.

After allele depth estimation, only 10% of markers were retained for the next analysis, in which we prioritized a minimum average allele depth of 60 reads to suppress the probable overestimation of allele bias due to our population size. Gerard et al. (2018) recommend that more than 25 reads be used to obtain a strong correlation of true genotypes under high levels of bias and overdispersion, emphasizing that read depth requirements should be based on how many individuals are included in the study. Autotetraploid linkage maps with higher-dose markers have recently been implemented in studies of tubers, such as autotetraploid potatoes (Massa et al., 2018; Mengist et al., 2018). In relation to mapping studies involving tropical forage grasses, only one reported the use of SNP markers with allele dosage, in which a minimum overall depth of 25 reads was considered in a biparental progeny of *U. decumbens* containing 217 F_1_ hybrids (Ferreira et al., 2019).

Our linkage map is the second for *M. maximus* but the first to be integrated from both parents and containing SNP markers in multiple doses. Updated statistical models and the recent technologies of genome sequencing promoted advances in the knowledge about the genetics of polyploid organisms. This is demonstrated by the comparison of our genetic map for *M. maximus* with the first map for this species published over ten years ago by Ebina et al. (2005). The first map contained 360 dominant markers obtained with amplified fragment length polymorphism (AFLP) and random amplified polymorphic DNA (RAPD) techniques that segregated at a 1:1 ratio and was obtained from an apomictic cultivar used for genetic breeding in Japan. These markers were distributed in 39 linkage groups, which is greater than the number expected for this species (2n = 4x = 32). In this context, the use of SNPs as codominant markers and their quantitative analysis allowed many gains in our map, such as coverage of many regions of quantitative traits important to breeders and the genetic effects that influence these traits, as well as alleles of the parents that were determinant of the characters of the hybrid.

In addition, the strategy of greater refinement of SNPs adopted after estimating allele dosage using two programs of mapping prevented overinflation among loci. We detected some markers with SD (Table 2), but we chose not to use them, aiming to increase the probability of obtaining an exact distance between markers in the HGs. Since genotyping errors are probable, a large amount of missing data and a large number of distorted loci may promote the expansion of linkage maps as well as overestimate the recombination fractions and limit the accuracy of the mapping (Cartwright et al., 2007; Gerard et al., 2018). However, SD is a phenomenon commonly found in the genome, and some linkage maps contain distorted markers, including those of grasses such as pearl millet (Sehgal et al., 2012) and napiergrass (Paudel et al., 2018). Therefore, greater knowledge of the occurrence and genetic causes of SD in plants is important for inferring which genes are kept together or separated by SD (Zhu et al., 2006; Anhalt et al., 2008). Thus, future studies may aggregate information on the structure of loci with SD in the genome of *M. maximus*.

Most linkage maps in grasses include only single-dose markers with segregation ratios of 1:1 and 3:1; thus, the heterozygous classes were grouped with one of the homozygous classes, resulting in a loss of information (Bourke et al., 2018b). The use of linkage maps with simplex SNPs for most linkage groups and a high density of higher-dosage markers provides greater confidence in the modeling of allelic effects of QTLs (Hackett et al., 2014). The markers present in our map were mostly simplex, while higher-dosage markers comprised approximately 15% of the map, as shown in Table 2. Similar values were also verified in the genetic map of *U. decumbens* (Ferreira et al., 2019), probably due to the complexity of scoring and analyzing these types of markers. Our map presented a greater number of alleles exclusive to the apomictic parent than sexual parent, as observed in the previous map of *M. maximus* (Ebina et al., 2005) and in the genetic maps of other forage grasses (Worthington et al., 2016; Ferreira et al., 2019). This difference is probably due to the origin of the genotypes, ‘Mombaça’ and S10 have natural tetraploid genomes but S10 was obtained from a sexual x apomictic cross of an original diploid sexual plant that was duplicated with colchicine.

Our linkage map, with an average density of 1.13 markers/cM (Table 3), was sufficient for the identification of monogenic and polygenic traits. Therefore, our map will also be useful for the detection of other important characteristics in *M. maximus* and can contribute to the assembly of the genome of this species, as well as to studies about the biology and evolution of other phylogenetically closely related tropical forage grasses, such as those in the *Urochloa* and *Paspalum* genera.

### 4.2 Double reduction in guinea grass

We reported for the first time the occurrence of DR in *M. maximus*. Autotetraploids may undergo this type of segregation when in multivalent pairing, two pairs of chromatids pass to the same pole in anaphase I of meiosis (Haynes and Douches, 1993). The distribution of the markers with DR between the dosage types was proportional to the number of SNPs with each configuration. DR has been extensively studied using SNPs with dosage data in autotetraploid linkage maps in potato, since the software for building maps was created to analyze data of this species (Hackett et al., 2017; Bourke et al., 2018a). Approximately 6% of markers in potato have DR (Bourke et al., 2015), corroborating our results (9.68%). Despite some studies suggesting that DR should be included in genetic map construction and in QTL analysis (Li et al., 2010), other studies verified that such markers have only minor positive effects on the power and accuracy of mapping analysis using single-dose markers (Bourke et al., 2015). However, no analyses have been performed for high-dose markers. Statistical models have been created to include DR in linkage mapping (Huang et al., 2019), but no software currently implements them.

The occurrence of DR in *M. maximus* has many implications for breeding programs, being that the effects of DR and how to handle them have been the targets of several studies (Luo et al., 2000; Xu et al., 2013; Layman and Busch, 2018). This type of segregation exposes alleles located in distal regions of the chromosomes to homozygosis and thus is effective in eliminating the lethal alleles in a population (Butruille and Boiteux, 2000). A low rate of DR is sufficient to considerably reduce the equilibrium frequency of a deleterious allele at one locus (Luo et al., 2006). As an alternative, DR could be used to accelerate the accumulation of favorable rare alleles through marker-assisted selection (MAS) (Bourke et al., 2015). In addition, it is possible to obtain genotypes with loci having a higher homozygosis rate for use in specific crosses (Bourke et al., 2015). In this context, more detailed molecular study could elucidate the influence of DR on the phenotypes of hybrids of our study species.

### 4.3 Apospory mapping and the search for gene similarity

A chi-square test (X^2^ = 5.43, p ≥ 0.01) performed for qualitative analysis of the reproductive mode of the 106 hybrids followed the Mendelian inheritance model, corroborating the results obtained with progeny tests in guinea grass performed by Savidan (1980) and Savidan (1981), in which sexual x apomictic progenies exhibited a 1:1 ratio that could be explained by an Aaaa genotype for apomicts, since the apomixis of *M. maximus* is dominant over sexuality. We could also prove this through SNP markers. Other apomixis studies in grasses such as *Pennisetum* (Akiyama et al., 2011), *Paspalum* (Martínez et al., 2003) and *Urochloa* (Valle et al., 1994; Vigna et al., 2016) also verified this segregation. Evidence suggests that this locus is present in a conserved region of the plant genome; however, further molecular genomic studies on apomixis in forage grasses are needed, since recent research has led to other hypotheses, such as a possible influence of epigenetics (Kumar, 2017).

The apomixis region in *M. maximus* was previously mapped (Ebina et al., 2005; Bluma-Marques et al., 2014), and similar to our results, no markers were in perfect linkage with the region. Nonetheless, we mapped markers at a shorter distance (0.8 cM) from the apo-locus (Figure 3). Genetic markers linked to apomixis have been sought in other forage grasses (Vigna et al., 2016; Worthington et al., 2016, 2019), aiming at the efficient and rapid identification of the reproductive mode of progenies. Once identified, such markers may be transferred among forage grasses, based on evidence of conservation of the ASGR. Markers near the apomixis region that were identified in our map may be validated and useful for the breeding program of this species.

In addition, we observed that the markers closer to the peak region of the apo-locus were in a genomic region of *P. virgatum* similar to a region of the genome of *A. thaliana* that contains the SERK1 gene. This gene is involved in the signaling pathway active during zygotic and somatic embryogenesis in *A. thaliana*, and its overexpression increases the efficiency of somatic embryogenesis initiation (Hecht et al., 2001). In grasses, studies about the genes involved in the regulation of reproductive events are relatively scarce. In nucellar cells of apomictic genotypes of *Poa pratensis*, the SERK gene is involved in embryo sac development (Albertini et al., 2005). Recently, SERK was reported in *Brachypodium distachyon* grass as having a domain conserved among monocots and plays a prominent role in apomixis (Oliveira et al., 2017).

### 4.4 QTLs for agronomic and nutritional traits

Traits related to the productivity and quality of forage used for cattle fattening are categorized as complex traits, which are determined by both genetic and environmental factors. The transgressive segregation observed in these traits in *M. maximus* suggests quantitative and polygenic inheritance, consistent with the inheritance of non-Mendelian traits (Miles and Wayne, 2008; Holzman and Hulsey, 2017). For the characterization of the genetic architecture of such traits, a QTL mapping approach is required. In our study, QTL analysis of autotetraploid progeny was performed using interval mapping of markers with allele dosage. This same methodology was successfully applied in QTL mapping in signalgrass, another important forage grass (Ferreira et al., 2019). The multiple interval mapping (MIM) method was recently implemented in polyploids and is a new alternative for other mapping studies using data with allele dose information (Pereira et al., 2019). The mapping method used here considered only the peak with the largest effect as a QTL; it is worth mentioning that peaks were present near the peak QTL for all agronomic traits (Supplementary Figure 2).

QTLs associated with important agronomic traits were mapped in HGs I, III, VII and VIII, with the phenotypic variation explained ranging from 4.3% to 10.4% (Table 4). Because *M. maximus* is undergoing a domestication process, the crop can still be greatly improved by the selection of large-effect QTLs. Therefore, QTL qRC9, located at 75 cM in HG II, which explained 10.3% of the phenotypic variation in RC (Supplementary Figure 2, HG_2(B)) and had a predominant additive effect of the female progenitor S10, may be a candidate for the marker-assisted selection program of guinea grass.

We found more than one QTL for TDM, LDM and RC, again supporting the hypothesis of complex genetic control. Conversely, we found only one QTL for PLB in both parents. TDM and LDM showed high broad-sense heritability (< 0.5), followed by RC (0.3) and PLB (0.1), as shown in Table 1. Higher heritability values (< 0.85) for LDM and RC and a value for PLB above 0.4 were recently reported in *M. maximus* (Lara et al., 2019), using the generalized heritability formula (Cullis et al., 2006). For this species, a greater amount (g/plant) of TDM and LDM in the progenies has been associated with considerable heritability from the most productive parents (Braz et al., 2013, 2017). Matias et al. (2019) also verified the same pattern in interspecific hybrids from *Urochloa* spp., another genus adapted to tropical conditions. The intermediate heritability of RC and the QTLs associated with this trait that were detected in both parents resulted in hybrids with a good capacity for regrowth. This trait is also considered fundamental in forage grasses because it is directly related to the persistence of the forage after defoliation (Jank et al., 2011).

The negative correlation between PLB and SDM was expected (Figure 1), and Braz et al. (2017) verified the higher PLB values in the experiment of this progeny. PLB is related to plant structure, and plants with a high percentage of leaves are desirable because this trait is related to higher forage quality. A higher PLB was observed in the male parent, cv. Mombaça, which is often used as a check in experiments. Since its release in the 1990s, along with cv. Tanzania, cv. Mombaça has promoted pasture intensification in the country due to its very high productivity and forage quality (Jank et al., 2014). Progenies whose female parent is S10 generally also present good yield (Resende et al., 2004). Breeding programs target these traits in search of superior genotypes with greater foliar mass and a higher percentage of leaves due to the higher digestibility of leaves than of stems for animals. Thus, forage breeding is not restricted to the obtaining of more productive plants; it also contributes to greater efficiency in their transformation into animal production (do Valle et al., 2009).

Significant and positive correlations were observed among GM, TDM, SDM, LDM, and RC, but not PLB, and corroborated the positions of QTLs associated with agronomic traits in the linkage map (Figures 1 and 3). We detected qTDM3 and qRC9 in a common region in HG II; qGM1, qTDM4, qLDM5, and qSDM7 in the same region in HG V; and qGM2, qLDM6, and qRC10 in part of a common region in HG VI. Each region containing several QTLs for different traits suggests the occurrence of four QTL hotspots. PLB presented a negative correlation with other agronomic traits and a positive correlation with NDF, but these correlations were weaker than those of the other pairs of traits (Figure 1). In the linkage map, it was possible to verify that QTLs for PLB were not detected in any regions with other agronomic traits; however, such a QTL was detected in a similar region with qNDF_L13 and qNDF_S18 in HG IV (Figure 3), suggesting a fifth QTL hotspot.

Clustering of QTLs for genetically correlated traits in the same or adjacent regions of HGs in several organisms may be due to physical linkage, pleiotropy or natural selection for coadapted traits (Studer and Doebley 2011; Wu et al. 2015). We have taken the first step in the identification of loci that lead to these trait correlations and the degree to which these patterns affect productivity in *M. maximus*. QTLs co-located in the same region of HGs for agronomic traits have also been identified in some grasses (Fang et al., 2016; Sartie et al., 2018).

Pleiotropy occurs when a gene influences multiple traits; when agronomically important traits are positively associated, more than one trait can be improved simultaneously (Kumar et al., 2017). There are some reports of QTLs with pleiotropic effects in some grasses, such as wheat (Deng et al., 2011), *Setaria* spp. (Mauro-Herrera and Doust, 2016), and a perennial grass (Khasanova et al., 2019).

Linked QTLs for natural selection of coadapted traits can occurs in two forms. First, these QTLs can be linked in the coupling phase on the chromosome, i.e., two favorable alleles of both genes are from the same parent, which is useful in breeding programs because can consider two traits as one target. Second, these QTLs are linked in the repulsion phase on the chromosome, where the two alleles of both genes are on opposite chromosomes, possibly due to favorable alleles of one gene being from one parent and the favorable alleles of another gene being from another parent. In this second case, linkage drag may occur, which needs to be broken for the breeding program (Wu et al., 2015).

All HGs contained QTLs related to nutritional traits, with those related to the leaf and stem being found mainly in HG III and HG VIII, respectively. Again, some traits had more than one peak that could not be considered (Supplementary Figure 3). The phenotypic variance explained by these QTLs ranged between 2.5% (qADF_L22) and 12.1% (qNDF_L11 and qNDF_S16). Interestingly, both parents contributed alleles for most of the identified QTLs, providing evidence that the genotypes have high nutritional quality.

The heritability of the nutritional traits varied from low (0.06), obtained for SIL_S, to moderate (0.32), obtained for OM_L. Traits IVD_L and IVD_S presented the same standard as the crude protein, with the H^2^ of the stem being higher than that of the leaf (Table 1). Historically, cv. Mombaça has stood out due to its high productivity, but with slightly lower values for forage quality when compared to ‘Tanzania’, and in biparental crosses, the hybrids obtained from ‘Mombaça’ also presented these features (Braz et al., 2017). This finding was corroborated by the heritability verified above in our study of agronomic and nutritional traits.

Notably, in the progenies of *M. maximus* from lower-yielding parents, the nutritional value is generally higher. With the identified QTLs, more in-depth studies of this correlation will be possible. In addition, QTLs related to forage quality have not been found in other important tropical species, such as *Urochloa* spp. Therefore, our results can contribute to the search for important genomic regions in other forage species. In contrast, in forage grasses of temperate climates, such as *Panicum hallii* (Milano et al., 2018) and *Lolium perenne* (Cogan et al., 2005), QTLs for nutritional and agronomic traits have been identified, even by using only single-dose markers.

The nutritional quality of forage grasses is important in several respects and is directly related to the production of meat and milk. *Megathyrsus maximus* shows a high value of CP compared to other tropical forages. However, the values for lignin and fiber are expected to be low because lignin hampers the enzymatic hydrolysis of cellulose and hemicellulose and, thus, the digestion of the cell wall of the leaf tissue and the stem (Jung and Allen, 1995). The relationship between the biomasses of leaves and stems is important due to its effects on nutritional value and voluntary consumption by animals. The NDF is associated with fibrous fractions and with voluntary consumption. The fractions that are not digested by the animal take up space in the digestive tract, impairing the digestion and consumption of dry matter (Euclides et al., 1999).

Complex correlations between nutritional traits were shown (Figure 1), especially that stem-related traits were most tightly correlated with leaf-related traits; however, NDF_L and NDF_S were closely correlated, and their respective QTLs were in the same regions in the HGs. The ADF, CEL and PL traits had a significant and positive correlation. In the linkage map, qCEL_L30 and qADF_L20 QTLs were in the same region, and qCEL_S31 and qADF_S26 were both located in HG VIII. Traits CP and IVD showed a weak positive correlation, but the qCP_S8 and qIVD_S28 QTLs were present in the same region of HG V, and qCP_S9 and qIVD_S29 were identified in the same region of HG VIII. The same pattern was observed for the leaf and stem, in which CP and IVD had a strong negative correlation with PL, ADF, CEL and NDF. Interestingly, qPL_S34 was verified in HG VIII in the same region as CP and IVD QTLs, and qADF_L21 shared a similar region with qCP_L7 in HG V. In addition, qCP_L7 extended to QTLs related to agronomic traits (qGM1, qTDM4, qLDM5 and qSDM7). A total of 8 probable QTL hotspots have been identified, supporting the need for further studies in search of a specific gene controlling all these traits or several genes acting together.

### 4.5 The search for similarity in apomixis and QTL regions

The search for putative candidate genes was based on all 23 QTL regions. Generally, the same gene families from *A. thaliana*, *O. sativa*, and *P. virgatum* were identified for a common QTL region. These genes are also found in the literature and are described in Table 6 and Supplementary Table 5.

Exploration of genes involved in plant growth and development, especially those related to hormone regulation, is crucial in forage grass breeding programs. Interestingly, the gibberellin family (GAS) was identified in HG III - region 9 (qPL_L33), whose QTL is related to lignin, a complex phenolic polymer deposited in the secondary cell wall of all vascular plants (Zhao, 2016). The interrelations between cell wall components cause cellulose and lignin to be codependent, a normal cellulose deposition pattern may be necessary for lignin assembly, and alterations of lignin content may lead to changes in the cell orientation of cellulose fibrils and, consequently, in digestibility (Anderson et al., 2015; Liu et al., 2016). GAs promote biochemical, physiological and anatomical plant changes (Hedden and Thomas, 2016). The induction of cellulose synthesis by GAs promotes the release of secondary regulators of cell wall proteins and, consequently, can boost lignin deposition and increase lignin content (Zhao, 2016). GAS at increased light levels have been shown to promote cell wall thickness and increase lignin deposition in xylem fibers (Falcioni et al., 2018).

Other important genes identified are associated with pectinesterase/pectin methylesterase and were present in HG V – region 14 (qGM1/qTDM4/qLDM5/qSDM7/qADF_L21/qCP_L7) and HG VIII – region 23 (qSIL_S37), which also contained agronomic QTLs. Pectinesterase is responsible for the hydrolyzation of pectin, the major component of cell walls (Pelloux et al., 2007). In addition, this enzyme is involved in developmental processes such as stem elongation in *A. thaliana* (Damm et al., 2015) and in *B. distachyon* grass (Feng et al., 2015). However, more in-depth research should be performed to ensure an understanding of the signaling pathways of these genes and make this understanding applicable to tropical forage breeding programs.

In conclusion, the present study produced a high-resolution linkage map with allele dosage information obtained from a full-sib progeny of *M. maximus* with high genetic variability. Even without the availability of a sequenced genome for this species, the approach adopted for the construction of our map was sufficient to detect many QTLs associated with agronomic and nutritional traits that are important for forage breeding. Our genetic map also allowed us to map the apo-locus to a single linkage group and provided a more up-to-date study of the mode of reproduction of *M. maximus*. The knowledge about the genetics of these traits that we obtained is the first step in discovering genes involved in relevant biological processes as well as understanding the genetic architecture of relevant traits in this species.

## Author contributions

AG, LJ and AdS conceived and designed the experiments. MS and LJ conducted the field experiments. TD, RF, AM, AP, and FO performed the laboratory experiments. TD, RF, LL, AM, AP, and FO analyzed the data. TD, RF and LL wrote the manuscript. All authors read and approved the manuscript.

## Acknowledgments

We acknowledge the members of the Brazilian Agricultural Research Corporation (Embrapa Beef Cattle) for making the progeny of this study available and for carrying out all phenotypic evaluations.

## Supplementary Material

**Figure S1.**
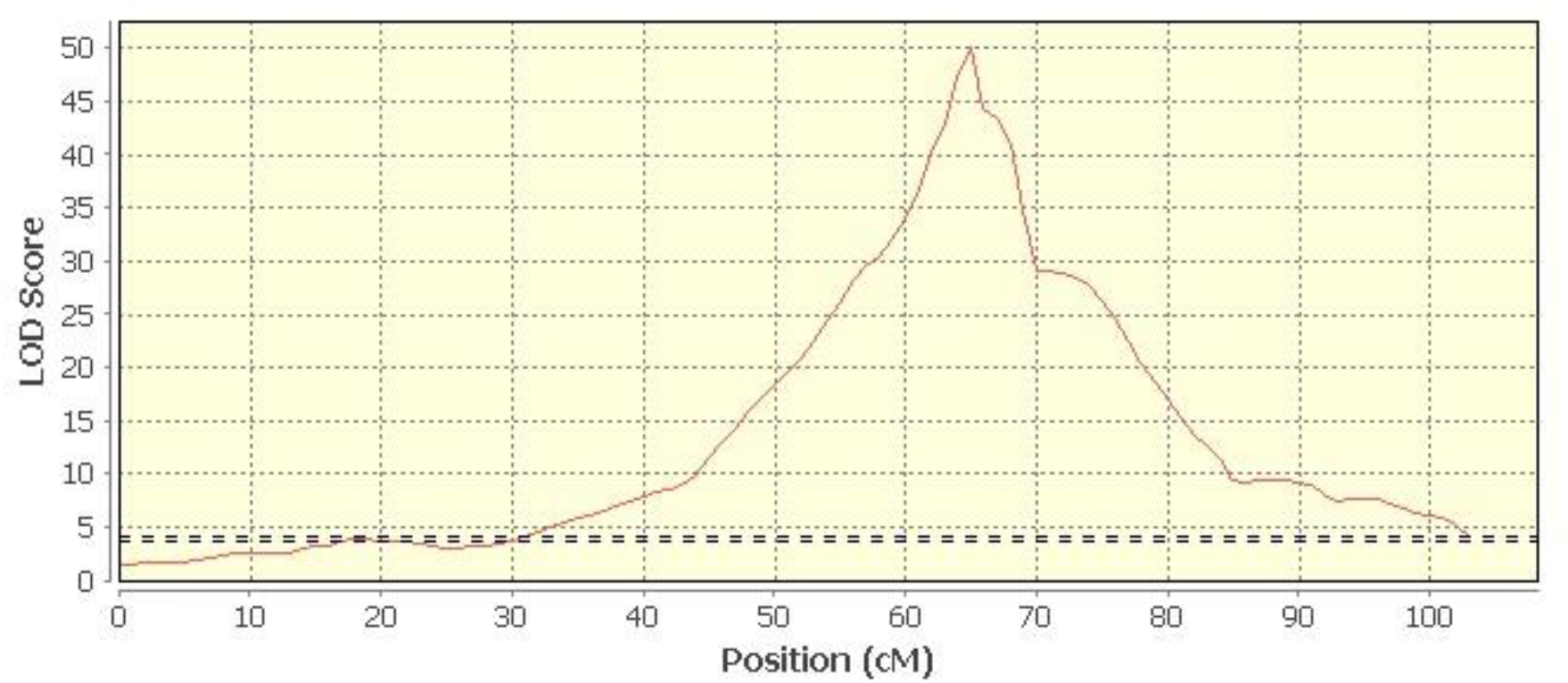
Identification of the apomixis region located in HG II from guinea grass (*Megathyrsus maximus*) mapping population. Dotted line indicate the LOD thresholds of 95% obtained after the permutation tests.

**Figure S2.**
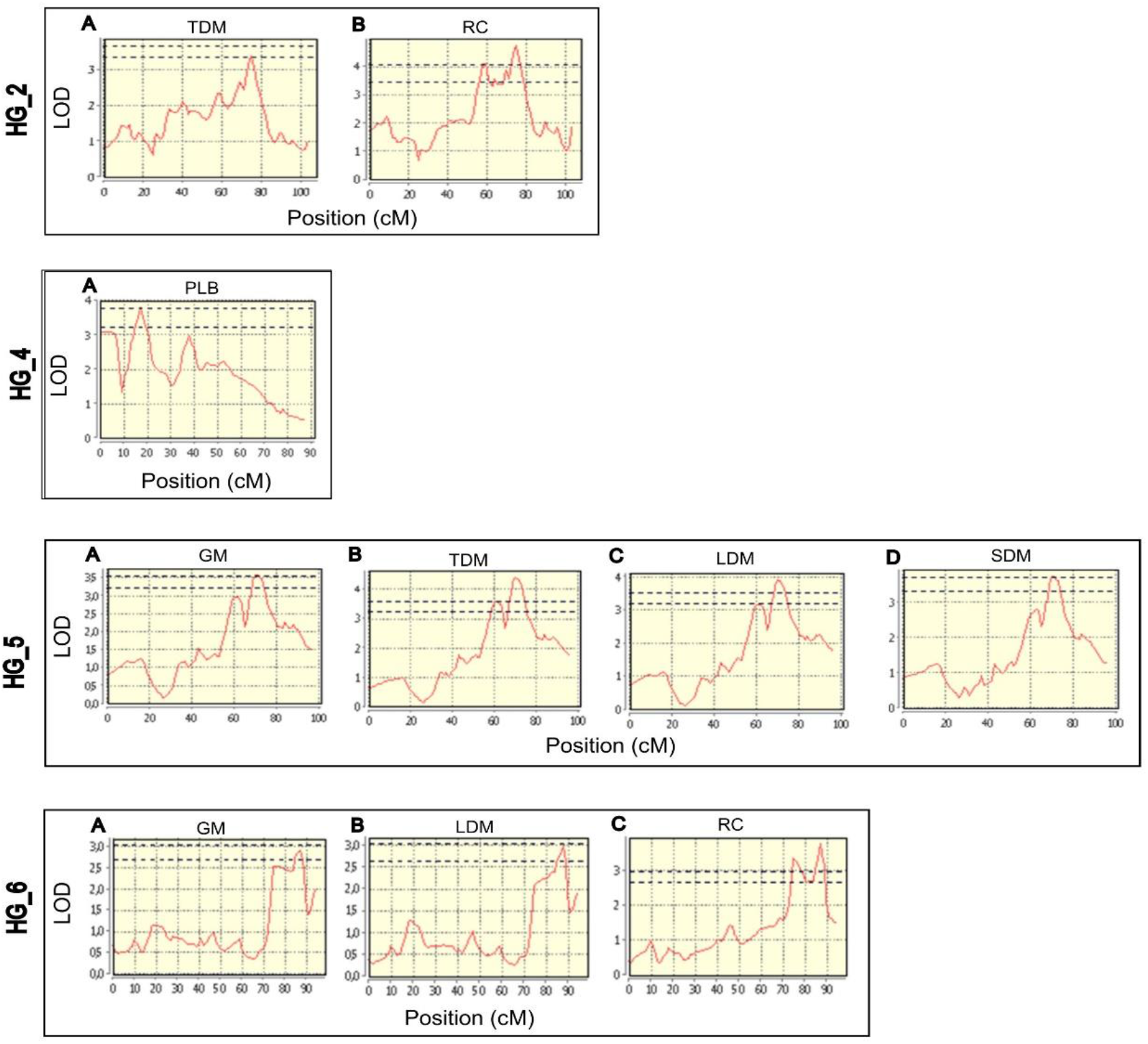
Interval mapping for agronomic traits from the guinea grass (*Megathyrsus maximus*) mapping population in HGs II, IV, V and VI.

**Figure S3(A).**
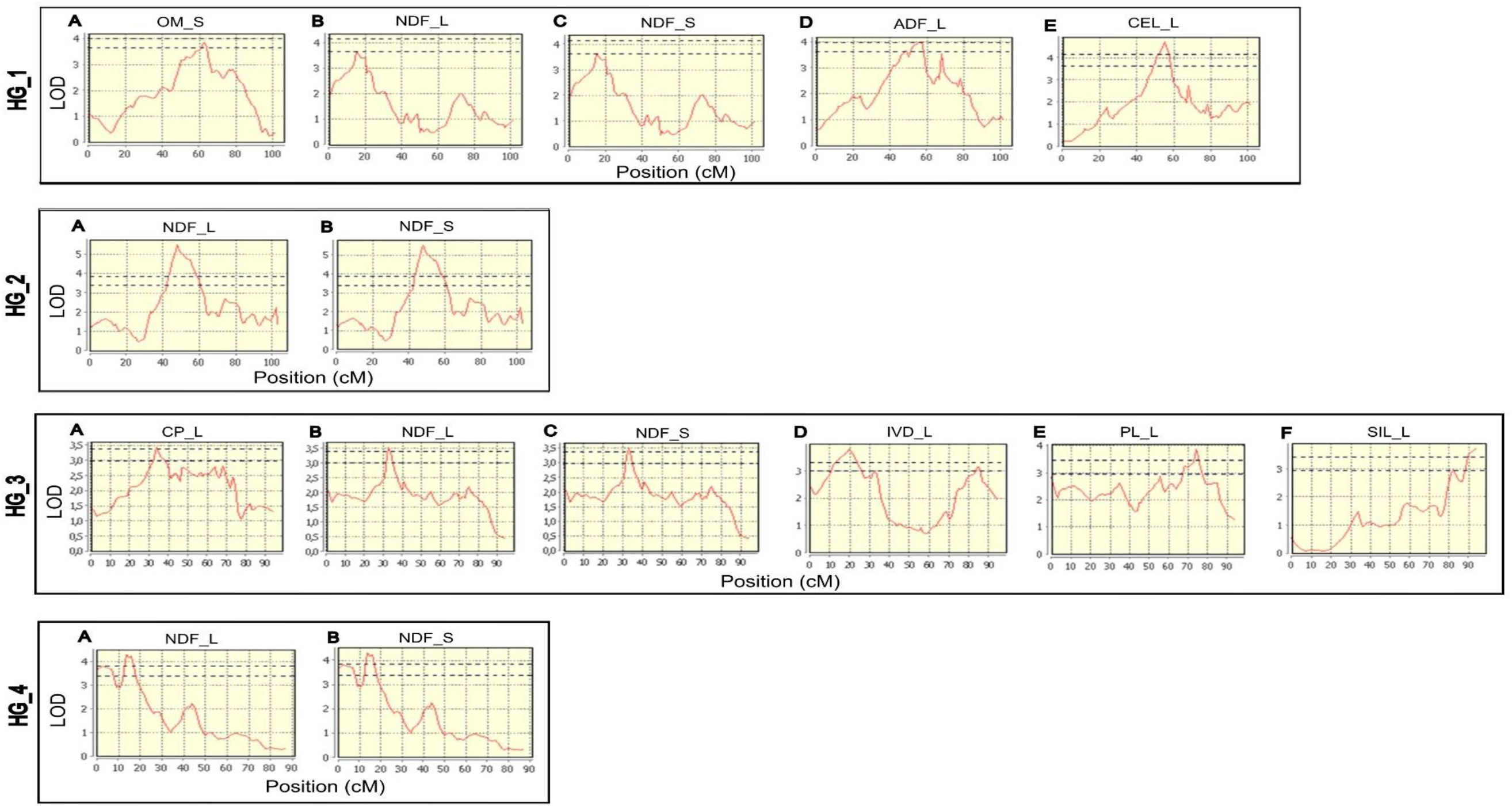
Interval mapping for forage quality from the guinea grass (*Megathyrsus maximus*) mapping population in HGs I to IV.

**Figure S3(B).**
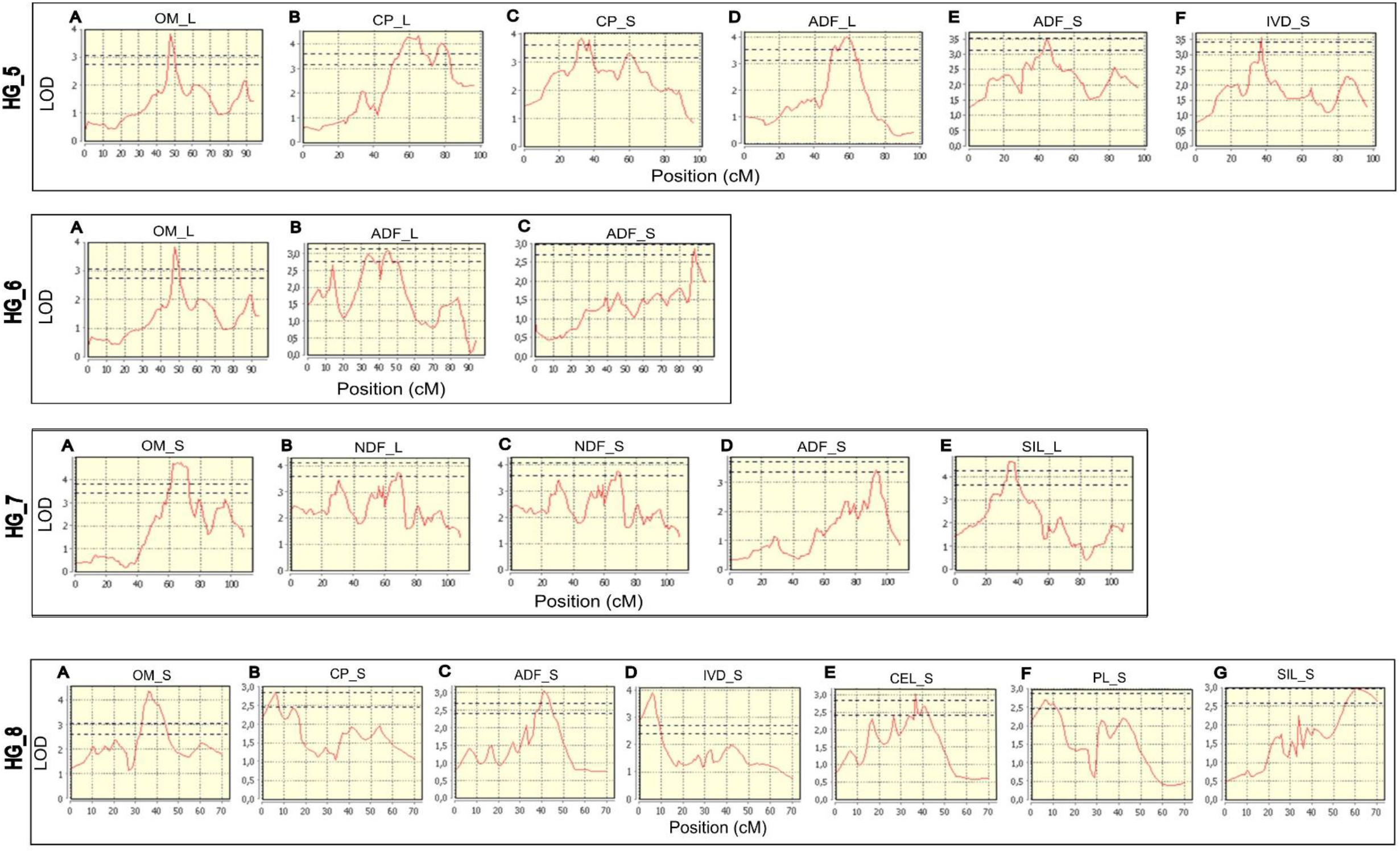
Interval mapping for forage quality from the guinea grass (*Megathyrsus maximus*) mapping population in HGs V to VIII.

**Table S1.** Reproductive mode of the hybrids from mapping population of guinea grass (*Megathyrsus maximus*).

| Genotype | RM<sup>a</sup> | Genotype | RM | Genotype | RM | Genotype | RM |
| --- | --- | --- | --- | --- | --- | --- | --- |
| B01 | Apo<sup>b</sup> | B46 | Apo | B102 | Sex | C24 | Apo |
| B02 | Sex<sup>c</sup> | B47 | Apo | B103 | Sex | C25 | Apo |
| B03 | ND<sup>d</sup> | B48 | Apo | B105 | ND | C26 | Apo |
| B04 | ND | B49 | Sex | B108 | Apo | C27 | Sex |
| B06 | Apo | B50 | Sex | B109 | Apo | C28 | Sex |
| B07 | ND | B51 | Apo | B110 | Sex | C30 | Sex |
| B10 | ND | B66 | Apo | B112 | ND | C31 | ND |
| B11 | Apo | B67 | Sex | B113 | Sex | C33 | Apo |
| B12 | Sex | B68 | Apo | B114 | Apo | C34 | Apo |
| B13 | Sex | B70 | Apo | B115 | Apo | C36 | Apo |
| B15 | ND | B71 | ND | B117 | Sex | C37 | ND |
| B16 | ND | B72 | Sex | B118 | Sex | C40 | Apo |
| B18 | Apo | B73 | Apo | B119 | ND | C41 | Apo |
| B20 | Sex | B74 | Sex | B120 | Apo | C43 | ND |
| B21 | Sex | B75 | Sex | B121 | ND | C44 | Apo |
| B22 | Sex | B76 | Apo | B124 | Apo | C45 | Sex |
| B23 | Sex | B77 | ND | B126 | Apo | C47 | Sex |
| B24 | Apo | B80 | ND | B128 | Apo | C48 | Sex |
| B25 | Sex | B81 | Sex | C02 | Apo | C50 | ND |
| B27 | ND | B83 | ND | C03 | Apo | C52 | Apo |
| B28 | Apo | B84 | Sex | C04 | Sex | C53 | Apo |
| B31 | ND | B85 | Sex | C07 | Apo | C54 | Sex |
| B33 | Sex | B91 | Apo | C09 | Apo | C55 | Apo |
| B34 | Apo | B92 | Apo | C10 | Apo | C57 | Apo |
| B35 | Sex | B93 | Apo | C12 | Apo | C58 | Apo |
| B37 | Apo | B94 | Apo | C13 | Sex | C59 | ND |
| B38 | Apo | B95 | ND | C14 | Apo | C60 | Apo |
| B39 | ND | B96 | Sex | C15 | Apo | C62 | Sex |
| B40 | Apo | B97 | Apo | C16 | Sex | C63 | Apo |
| B41 | Apo | B98 | ND | C17 | Sex | C64 | Apo |
| B42 | ND | B99 | Apo | C21 | Apo | C65 | Sex |
| B44 | Apo | B100 | Apo | C22 | Sex | C66 | Sex |
| B45 | Apo | B101 | Apo | C23 | Apo | C90 | ND |

<sup>a</sup>Reproductive mode; <sup>b</sup>Apomictic; <sup>c</sup>Sexual; <sup>d</sup>Not determined.

**Table S2.** AIC and SIC values for ***G_L_*** and ***R_L_*** matrices for agronomic traits. The lowest values and selected variance-covariance (VCOV) structures are indicated in bold. NC means not converged.

| GM |  |  |  |  |  | TDM |  |  |  |  |  | LDM |  |  |  |  |  |
| --- | --- | --- | --- | --- | --- | --- | --- | --- | --- | --- | --- | --- | --- | --- | --- | --- | --- |
| $G_L$ matrix | | | $R_L$ matrix | | | $G_L$ matrix | | | $R_L$ matrix | | | $G_L$ matrix | | | $R_L$ matrix | | |
| VCOV | AIC | SIC | VCOV | AIC | SIC | VCOV | AIC | SIC | VCOV | AIC | SIC | VCOV | AIC | SIC | VCOV | AIC | SIC |
| ID | 1610.4 | 1632.8 | ID | 581.7 | 598.5 | ID | 1730.0 | 1752.5 | ID | 864.3 | 886.7 | ID | 1738.9 | 1761.3 | ID | 823.5 | 840.3 |
| DIAG | 1619.6 | 1670.2 | DIAG | 569.4 | 614.4 | DIAG | 1738.2 | 1788.7 | DIAG | 836.7 | 887.2 | DIAG | 1747.5 | 1798.0 | DIAG | 798.6 | 843.5 |
| CS | 581.7 | **598.5** | CS | -272.8 | -239.1 | CS | 871.8 | 888.7 | CS | 216.8 | 250.5 | **CS** | 823.5 | **840.3** | CS | 87.4 | 115.5 |
| CSHet | 587.4 | 632.4 | CSHet | -295.7 | -233.9 | CSHet | 871.0 | 915.8 | CSHet | 170.4 | 232.1 | CSHet | 828.6 | 879.1 | CSHet | 46.4 | 108.1 |
| AR1 | NC | NC | AR1 | -546.0 | -512.3 | **AR1** | **864.3** | **886.7** | AR1 | 241.7 | 275.3 | AR1 | **818.8** | 846.9 | AR1 | NC | NC |
| AR1Het | NC | NC | AR1Het | -568.0 | -506.2 | AR1Het | NC | NC | AR1Het | 168.9 | 230.6 | AR1Het | NC | NC | **AR1Het** | **-92.1** | **-30.4** |
| FA1 | 585.3 | 635.8 | FA1 | NC | NC | FA1 | 870.0 | 926.1 | FA1 | NC | NC | FA1 | NC | NC | FA1 | NC | NC |
| US | **567.3** | 702.2 | **US** | **-713.8** | **-573.3** | US | 873.9 | 1003.0 | **US** | **-9.5** | **130.8** | US | NC | NC | US | NC | NC |
| SDM |  |  |  |  |  | PLB |  |  |  |  |  | RC |  |  |  |  |  |
| $G_L$ matrix | | | $R_L$ matrix | | | $G_L$ matrix | | | $R_L$ matrix | | | $G_L$ matrix | | | | | |
| VCOV | AIC | SIC | VCOV | AIC | SIC |  | AIC | SIC |  | AIC | SIC |  | AIC | SIC |  |  |  |
| ID | 965.4 | 980.1 | ID | 835.8 | 850.5 | ID | 1954.4 | 1971.3 | ID | 1831.0 | **1861.4** | ID | 377.2 | 421.6 |  |  |  |
| DIAG | 966.4 | 990.9 | DIAG | 824.8 | 854.3 | DIAG | 1950.1 | 1995.0 | **DIAG** | **1821.8** | 1872.3 | DIAG | 378.1 | 427.5 |  |  |  |
| CS | 835.8 | **850.5** | CS | 806.6 | **831.2** | CS | 1848.6 | 1871.0 | CS | 1839.5 | 1867.6 | **CS** | **343.0** | **387.3** |  |  |  |
| CSHet | **834.3** | 858.8 | CSHet | 798.0 | 837.4 | CSHet | 1836.1 | 1886.6 | CSHet | 1822.4 | 1878.5 | CSHet | 345.7 | 400.0 |  |  |  |
| AR1 | 836.0 | 850.8 | AR1 | 826.0 | 845.7 | **AR1** | 1839.0 | **1861.4** | AR1 | 1840.8 | 1868.8 | AR1 | 344.9 | 394.2 |  |  |  |
| AR1Het | 834.4 | 859.0 | AR1Het | 815.4 | 849.8 | AR1Het | **1827.5** | 1878.0 | AR1Het | 1823.7 | 1879.9 | AR1Het | 345.7 | 400.0 |  |  |  |
| FA1 | 836.1 | 865.7 | FA1 | NC | NC | FA1 | 1838.1 | 1911.1 | FA1 | NC | NC | FA1 | 346.8 | 406.0 |  |  |  |
| US | 840.3 | 879.6 | **US** | **793.3** | 837.6 | US | NC | NC | US | 1833.3 | 1968.0 | US | 352.1 | 421.1 |  |  |  |

**Table S3.** AIC and SIC values for ***G_L_*** and ***R_L_*** matrices for nutritional traits.

| Leaf |  |  |  |  |  |  |  |  |
| --- | --- | --- | --- | --- | --- | --- | --- | --- |
|  | OM_L | CP_L | NDF_L | ADF_L | IVD_L | CEL_L | PL_L | SIL_L |
| AIC | 342.34 | 500.52 | 354.96 | 345.76 | 329.68 | 528.47 | 348.05 | 343.99 |
| SIC | 353.84 | 515.86 | 362.62 | 357.26 | 341.19 | 536.14 | 359.55 | 355.49 |
| Stem |  |  |  |  |  |  |  |  |
|  | OM_S | CP_S | NDF_S | ADF_S | IVD_S | CEL_S | PL_S | SIL_S |
| AIC | 328.04 | 310.15 | 1019.01 | 666.55 | 332.18 | 467.56 | 319.97 | 337.44 |
| SIC | 339.44 | 321.55 | 1026.67 | 677.95 | 343.58 | 475.16 | 331.37 | 348.83 |

**Table S4**. Name, position and dosage type of the SNP markers in the eight HGs.

**Table S5**. Putative candidate genes identified in apomixis and QTL regions from *P. virgatum*, *A. thaliana* and *O. sativa*.

